# HCV spread kinetics reveal varying contributions of transmission modes to infection dynamics

**DOI:** 10.1101/2021.04.12.439579

**Authors:** Karina Durso-Cain, Peter Kumberger, Yannik Schälte, Theresa Fink, Harel Dahari, Jan Hasenauer, Susan L. Uprichard, Frederik Graw

**Affiliations:** Department of Microbiology and Immunology and Infectious Disease and Immunology Research Institute, Stritch School of Medicine, Loyola University Medical Center, Maywood, Illinois, USA; The Program for Experimental and Theoretical Modeling, Division of Hepatology, Department of Medicine, Stritch School of Medicine, Loyola University Medical Center, Maywood, Illinois, USA; BioQuant-Center for Quantitative Biology, BIOMS, Heidelberg University, 69120 Heidelberg, Germany; Mathematical Modeling of Biological Systems, Department of Mathematics, TU München, 85748 Garching, Germany; Institute of Computational Biology, Helmholtz Zentrum München, 85764 Neuherberg, Germany; Faculty of Mathematics and Natural Sciences, Rheinische Friedrich-Wilhelms-Universität Bonn, 53115 Bonn, Germany

**Keywords:** HCV, cell-to-cell transmission, mathematical modeling, spatial spread, agent-based model

## Abstract

Hepatitis C virus (HCV) is capable of spreading within a host by two different transmission modes: cell-free and cell-to-cell. Although viral dissemination and diffusion of viral particles facilitates the infection of distant cells, direct cell-to-cell transmission to uninfected neighboring cells is thought to shield the virus from immune recognition. However, the contribution of each of these transmission mechanisms to HCV spread is unknown. To dissect the contribution of these different transmission modes to HCV spread, we measured HCV lifecycle kinetics and used an in vitro spread assay to monitor HCV spread kinetics after low multiplicity of infection in the absence and presence of a neutralizing antibody that blocks cell-free spread. By analyzing these data with a spatially-explicit mathematical model that describes viral spread on a single-cell level, we quantified the contribution of cell-free and cell-to-cell spread to the overall infection dynamics and show that both transmission modes act synergistically to enhance the spread of infection. Thus, the simultaneous occurrence of both transmission modes likely represents an advantage for HCV that may contribute to the efficient establishment of chronic infection. Notably, the relative contribution of each viral transmission mode appeared to vary dependent on different experimental conditions and suggests that viral spread is optimized according to the environment. Together, our analyses provide insight into the transmission dynamics of HCV and reveal how different transmission modes impact each other.

**Importance:** Hepatitis C Virus can spread within a host by diffusing viral particles or direct cell-to-cell transfer of viral material between infected and uninfected cells. To which extend these cell-free and cell-to-cell transmission modes contribute to HCV spread, establishment of chronicity and antiviral escape is still unknown. By combining *in vitro* experimental HCV spread data with a multi-scale mathematical model we have disentangled the contribution and interplay of cell-free and cell-to-cell transmission modes during HCV infection. Our analysis revealed synergistic effects between the two transmission modes, with the relative contribution of each transmission mode varying dependent on the experimental conditions. This highlights the adaptability of the virus and suggests that transmission modes might be optimized dependent on the environment, which could contribute to viral persistence.

## Introduction

The way a virus spreads within a host is a critical determinant that impacts the establishment and progression of an infection that can affect pathogenesis, host response and treatment efficacy. Although classical viral life cycles are often diagramed as being initiated by entry of diffusing virions via cell-surface receptors followed by viral replication and subsequent release of newly formed viral particles, it is recognized that viruses can spread by multiple ways. Besides transmission by cell-free viral particles, many viruses [1–3], including Hepatitis C virus (HCV) [4, 5], have been observed to spread via direct cell-to-cell mechanisms. The strategies employed by different viruses are not all well-defined, but can involve a broad range of mechanisms, such as the formation of virological synapses, movement on the outside of membrane bridges or extensions created by either the target or donor cell, or within cytoplasmic tunnels connecting adjacent cells [1, 2]. In terms of efficient dissemination of infection, both cell-free and cell-to-cell transmission have their advantages and disadvantages. While diffusing viral particles facilitate the infection of distant cells and transmission to new hosts, direct cell-to-cell transmission between neighboring cells is considered to be more efficient as it can circumvent complex entry processes and shield viral material from neutralizing antibodies [5–7]. Furthermore, it is thought to allow more viral genomes to simultaneously enter individual cells, increasing resistance to host defenses and antiviral therapies [8]. Yet, to which extent these different means of transmission contribute to viral spread and establishment of chronic infection, as well as influence viral escape and disease progression has not been determined.

Cell-free and cell-to-cell spread can be studied individually by experimentally blocking either of the two transmission modes. For HCV, neutralizing antibodies against the HCV E2 glycoprotein (anti-E2) have been used to inhibit cell-free virus uptake and to allow the study of HCV cell-to-cell transmission *in vitro* [4, 9–11]. However, when both transmission modes occur simultaneously, the extent to which each of these transmission modes contributes to viral spread and influences the other is difficult to measure directly. Mathematical models that provide an accurate representation of the infection process have proven to be invaluable for analyzing infection dynamics and quantifying important parameters that characterize viral spread kinetics and response to antiviral treatment (reviewed in [12]), particularly in the case of HCV [13–16]. Specifically, analyzing the interplay of cell-free and cell-to-cell transmission during viral spread requires the use of mathematical models that are able to account for the spatio-temporal dynamics of these processes and the individual dynamics of foci growth [12, 17].

In this study, we combined experimental data and mathematical modeling to determine the contribution and dynamic interaction between cell-free and cell-to-cell transmission modes in HCV spread. Experimentally, we analyzed the kinetics of HCV single cycle infection at the population level measuring intracellular replication and extracellular viral secretion, as well as the spatial spread of HCV during multi-cycle infection on a single cell level by measuring foci number and growth under different conditions, e.g. in the absence and presence of neutralizing antibodies that block cell-free transmission. To analyze these data, we developed a spatially resolved, multi-scale agent-based model (ABM) that accounts for intracellular viral replication dynamics, direct cell-to-cell spread, as well as extracellular viral diffusion and cell-free spread. Using appropriate parameter inference methods to adapt our multi-scale ABM to the experimental data, our model is able to describe the experimentally observed spread dynamics. The model allows for estimates of transmission parameters and to infer the contribution of cell-free and cell-to-cell transmission to HCV spread that cannot be measured experimentally. We found that the relative contribution of each transmission mode varied under different culture conditions, which suggests that HCV may optimize the spread mechanisms utilized according to the environmental conditions. Together, our analyses provide insight into the transmission dynamics of HCV on a cellular level and reveal how different transmission modes might impact each other.

## Results

### Single round infection kinetics defines the timing of viral life cycle

Viral spread can only occur after sufficient time has elapsed to allow for infectious progeny virus production in the newly infected cell. Therefore, to elucidate the kinetics of HCV spread, we determined the timing of different HCV life cycle events, by analyzing the kinetics of intracellular and extracellular viral RNA. To this end, Huh7 cells were infected with HCV at a multiplicity of infection (MOI) of 6 for 3 hours to establish a reasonably synchronized infection. Culture media and cell lysates were then harvested from triplicate wells at frequent time intervals for 96 hours. Total RNA was extracted from the cell lysates and the culture media to quantify intracellular and extracellular HCV RNA levels by RT-qPCR (**Fig. 1A** and **B**, respectively). The culture media was also titrated to determine extracellular infectious HCV levels (**Fig. 1B**). Intracellular HCV RNA first increased between 9 and 12 hours post-inoculation (p.i.), suggesting HCV RNA replication begins around 9 hours p.i. Intracellular RNA levels then continued to increase until approximately 72 hours p.i. when levels began to plateau (**Fig 1A**). After ~18 hours of decline, extracellular HCV RNA plateaued and then both extracellular RNA and viral titers started to slowly increase (**Fig. 1B**), which suggests that secretion of progeny virus begins at ~18 hours p.i., but that newly secreted virus levels are initially tempered by continued degradation of input virus and possibly some disappearance of extracellular virus due to infection. Based on the early kinetics observed, secondary cell-free virus infection would be expected to begin contributing to detected intracellular HCV RNA accumulation levels by 27 hours p.i. (i.e. 18 hours for secondary infection to initiate + 9 hours for replication to occur), and extracellular HCV RNA and titer levels by 36 hours p.i. (i.e. 18 hours for secondary infection to initiate +18 hours for secretion of progeny virus to begin). However, cell-to-cell spread could in theory begin sometime after 9 hours of infection as newly synthesized genomic HCV RNA is accumulating in the cell. Additionally, this data indicates that to experimentally inhibit cell-free spread, the necessary inhibitor (e.g. neutralizing HCV anti-E2) needs to be added prior to 18 hours p.i. before infectious progeny virus is detected.

**Figure 1:**
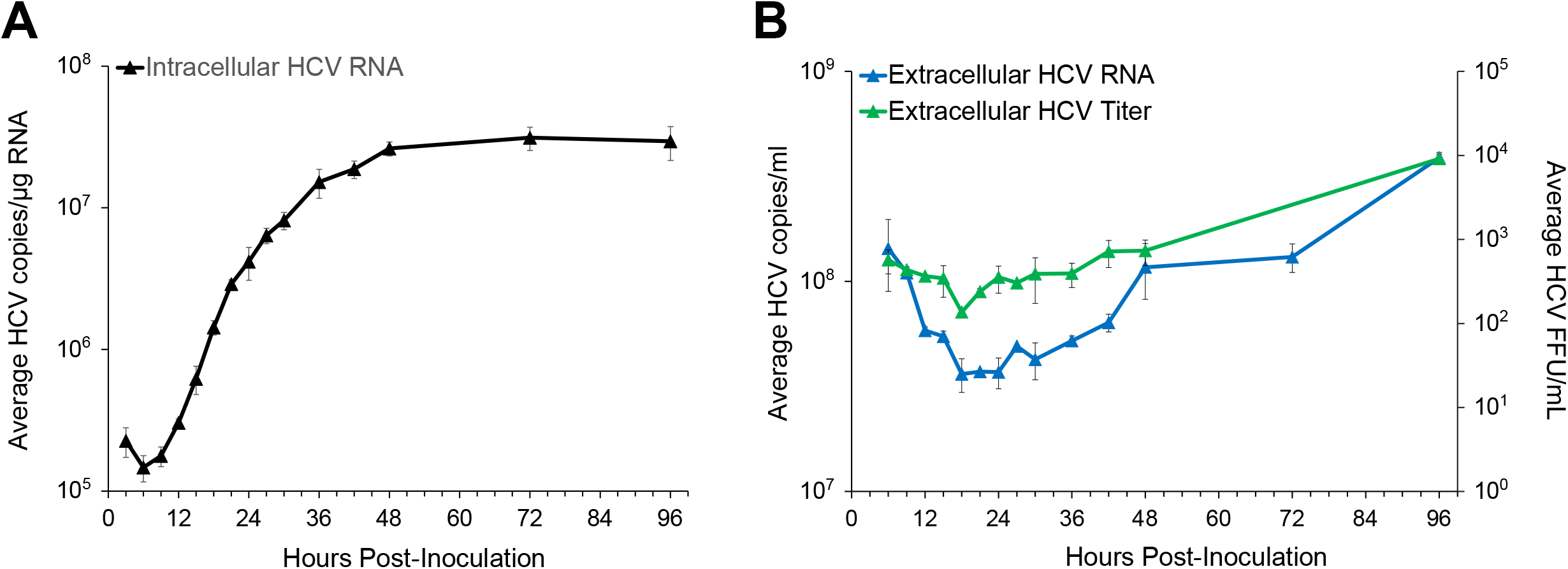
HCV High MOI Infection Kinetics: Huh7 cells were infected with HCV at a MOI of 6. Cell lysates and culture media was harvested at the indicated time points. **(A)** Intracellular HCV RNA (black triangles). HCV and cellular GAPDH RNA levels were quantified by RT-qPCR. Cellular GAPDH was used to normalize HCV copies, which are then graphed as average HCV copies/μg RNA in triplicate samples +/- standard deviation (SD). **(B)** Extracellular HCV RNA (blue triangles) are graphed as average HCV copies/mL. HCV titers (green triangles) are graphed as foci-forming units (FFU)/mL. Pooled culture media samples from the triplicate wells were spiked with equal amounts of mouse liver RNA as an internal control before extracellular RNA was extracted. Extracellular HCV RNA and mouse GAPDH (mGAPDH) RNA levels were quantified by RT-qPCR. HCV copies were normalized to mGAPDH and graphed as average HCV copies/mL in duplicate samples +/- SD. Titers were determined by titrating the pooled media on naïve Huh7 cells. Results are the average of foci counted in 3 wells +/- SD.

### Monitoring viral spread in the presence and absence of neutralizing antibodies that block cell-free infection

To characterize the impact of cell-to-cell transmission on HCV spread, we measured viral spread in untreated cultures where both cell-free and cell-to-cell spread occur together and used a neutralizing E2 antibody (anti-E2) in parallel cultures to inhibit cell-free spread and measure cell-to-cell transmission alone. For each of these conditions, HCV spread was analyzed by evaluating the number and size of HCV infected cell foci (**Fig. 2A**). In untreated wells where both cell-free and cell-to-cell spread were free to occur there was an increase in mean focus size over time (mean and [range]: 4.73 [1, 66] cells/focus at 72 hpi; 23.1 [1, 227] cells/focus at 120 hpi). However, there was also an increase in the number of small foci, which resulted in only a slight increase in the median focus size (2 cells/focus at 72 hpi vs. 4 cells/focus at 120 hpi) (**Fig. 2B**). Thus, focus sizes showed a bimodal distribution with many small and large foci at later time points (**Fig. 2B; Fig. S1**). In contrast, in cultures where anti-E2 was added to block cell-free spread, there was an increase in foci sizes over time without an increase in number of foci, and, thus, no establishment of new small foci, resulting in a steady increase in median focus size over time (4 cells/focus at 72 hpi vs. 18.5 cells/focus at 120 hpi) (**Fig. 2B**). Largest foci at 96 and 120 hours p.i. were observed in the untreated wells. Besides the possibility of foci merging, for which we did not find evidence in our observations, this appeared to suggest that cell-free virus spread was not only establishing new small foci, but also contributing to the growth of individual foci. The difference in the average total number of infected cells of 1560 +/-387 and 938 +/-158 infected cells at 120 hours p.i. in untreated and anti-E2 treated wells, respectively showed that wells in which cell-free spread was additionally able to occur exhibited a ~1.7-fold higher number of infected cells (**Fig. 2D**).

**Figure 2:**
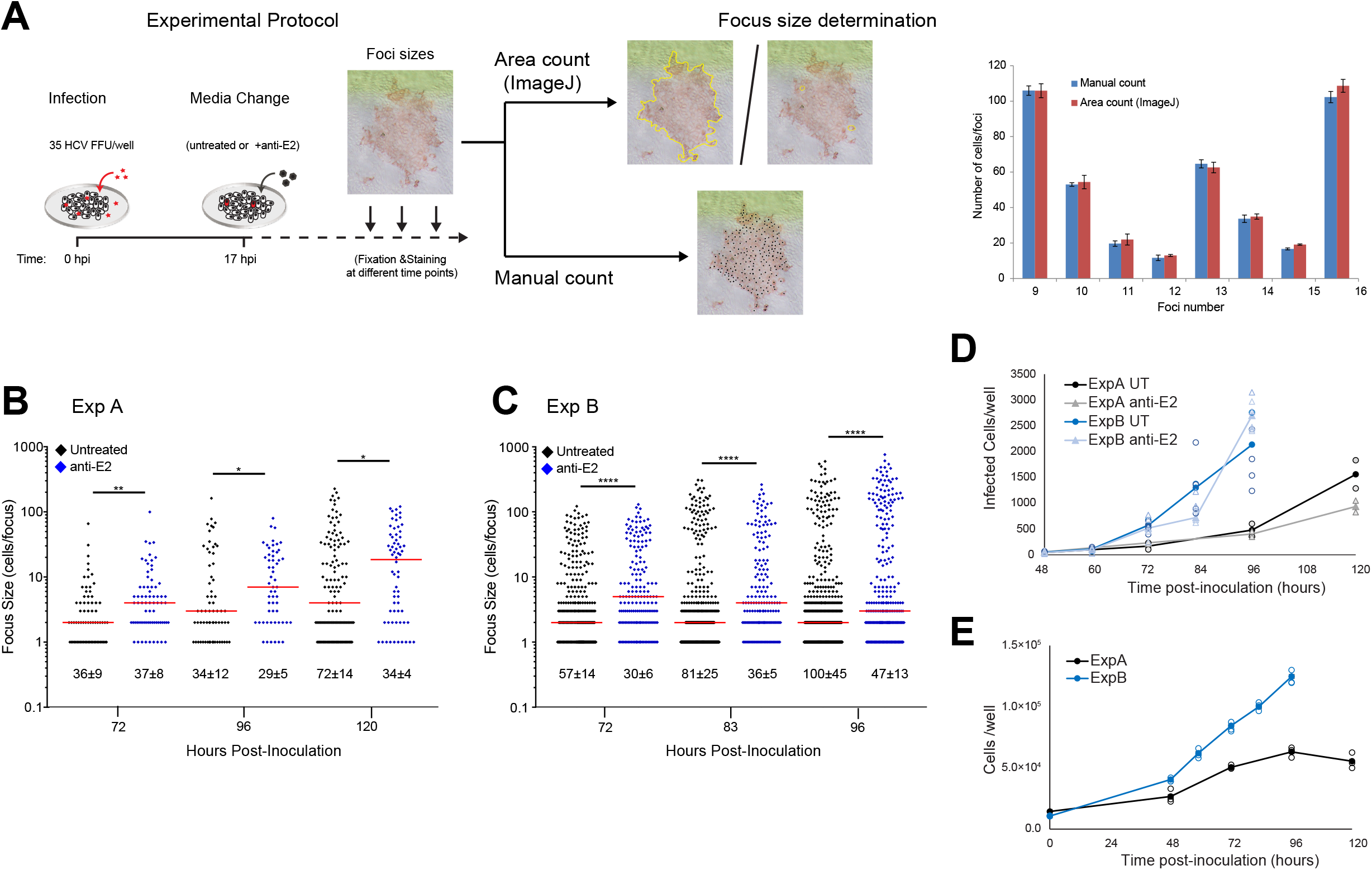
HCV Spread Kinetics: **(A)** Schematic of the experimental protocol: Confluent Huh7 cells were infected with 35 HCV FFU/well. At 17 hours p.i. the viral inoculum was removed and fresh cDMEM was added either in the absence (i.e. untreated) or presence of 10μg/mL anti-E2. Monolayers were fixed at different times post-infection and then stained for HCV to detect infected cells. Focus size was counted by individual cell counts for small foci or by area using ImageJ to circle each focus and divide the focus area by the average area of single cells for larger foci. The accuracy of ImageJ focus size calculation was determined by comparing 3 technical replicate ImageJ area counts with 3 technical replicate manual counts. Graph shows individual cell counts vs. area counts by ImageJ for representative foci, numbered arbitrarily on the X-axis. **(B)** Representative of early experiments: Duplicate wells fixed at 72, 96 and 120 hours p.i. and then stained for HCV with NS5A antibody to detect infected cells. **(C)** Representative of later experiments: Five to six wells were fixed at 72, 83 and 96 hours p.i. and stained for HCV with E2 antibody to detect infected cells. The numbers of foci per well were counted and are indicated below the dot plots as average +/- standard deviation per well. The number of cells/focus were counted and graphed. Red bar = Median focus size. Statistical differences relative to untreated are indicated (*, P<0.05; **, P<0.01; ****, P<0.0001 by Mann-Whitney U-test). Note that foci sizes from individual wells were pooled for the statistical comparison of foci sizes. The results in each panel are representative of two experimental repeats. **(D)** Total number of infected cells per well over time calculated from the data in **B** and **C**. Individual well counts (open circles/triangles) and mean (closed circles/triangles). **(E)** Total number of cells per well over time counted in parallel untreated wells for each experiment. Individual (open circles) and mean (closed circles) cell numbers for three wells are shown.

Surprisingly, a subsequent round of experiments gave a different result with median foci sizes in anti-E2 treated wells decreasing over time from 72 hours to 96 hours (**Fig. 2C**). One obvious explanation for these differences in focus growth patterns was viral escape from anti-E2 neutralization as evidenced by the increasing number of foci in the anti-E2 treated wells in the later experiments (**Fig. 2C**). While the presence of viral escape was perplexing considering that the identical protocol and the same pre-aliquoted cells, virus and antibody were utilized for all experiments, one difference identified was an increase in the number of cell divisions that occurred during the later experiments, which in hindsight correlated to the purchase of a new lot of fetal bovine serum (FBS). Specifically, the earlier round of experiments (represented in **Fig. 2B**, Exp. A) exhibited a 2.5-fold increase in cell numbers at 96 hours post plating, while the later experiments (represented in **Fig. 2B**, Exp. B) exhibited a ~12-fold increase in cell numbers 96 hours post plating (**Fig. 2E**) despite the same number of cells being plated at confluence for all experiments. We speculate that the increase in cell number allowed for higher virus levels per well which outcompeted the available neutralizing antibody and allowed for cell-free spread in anti-E2 treated wells. However, it remains to be determined to which extent cell-free and cell-to-cell transmission modes contribute to HCV spread within these different settings.

### A multi-scale mathematical model to analyze HCV spread kinetics

To further elucidate the dynamics of HCV spread and disentangle the contribution of cell-free and cell-to-cell transmission, we extended an agent-based model that we had developed previously for analyzing the spread of HCV *in vitro* [14, 17]. The multi-scale model follows the progression of infection on a single-cell level by considering individual cell infection kinetics as well as their spatial distribution in the monolayer of cells to account for local effects. It combines a deterministic description of the intracellular viral replication and secretion dynamics with the stochastic transmission dynamics on a cellular level in the form of an agent-based model with cells representing the individual agents. A sketch of the model is provided in **Figure 3A** with a detailed description provided in *Materials and Methods*. In brief, uninfected cells are distributed on a hexagonal grid representing the *in vitro* culture system. These cells proliferate and can get infected via either cell-free or cell-to-cell transmission according to stochastic processes that depend on the extracellular viral concentration at the respective grid site or the intracellular viral concentration of directly neighboring infected cells, respectively. Upon infection, the intracellular processes of viral replication and viral export for each individual cell are described by a deterministic mathematical model according to the viral kinetics observed experimentally. This model describes the logistic growth of intracellular RNA, *R*, according to a maximal production rate λ with viral RNA exported to the extracellular space at a constant rate ρ (**Fig. 3A** and *Materials and Methods)*. Extracellular viral RNA will diffuse through the simulated grid to contribute to cell-free transmissions and is cleared at rate *c*.

**Figure 3:**
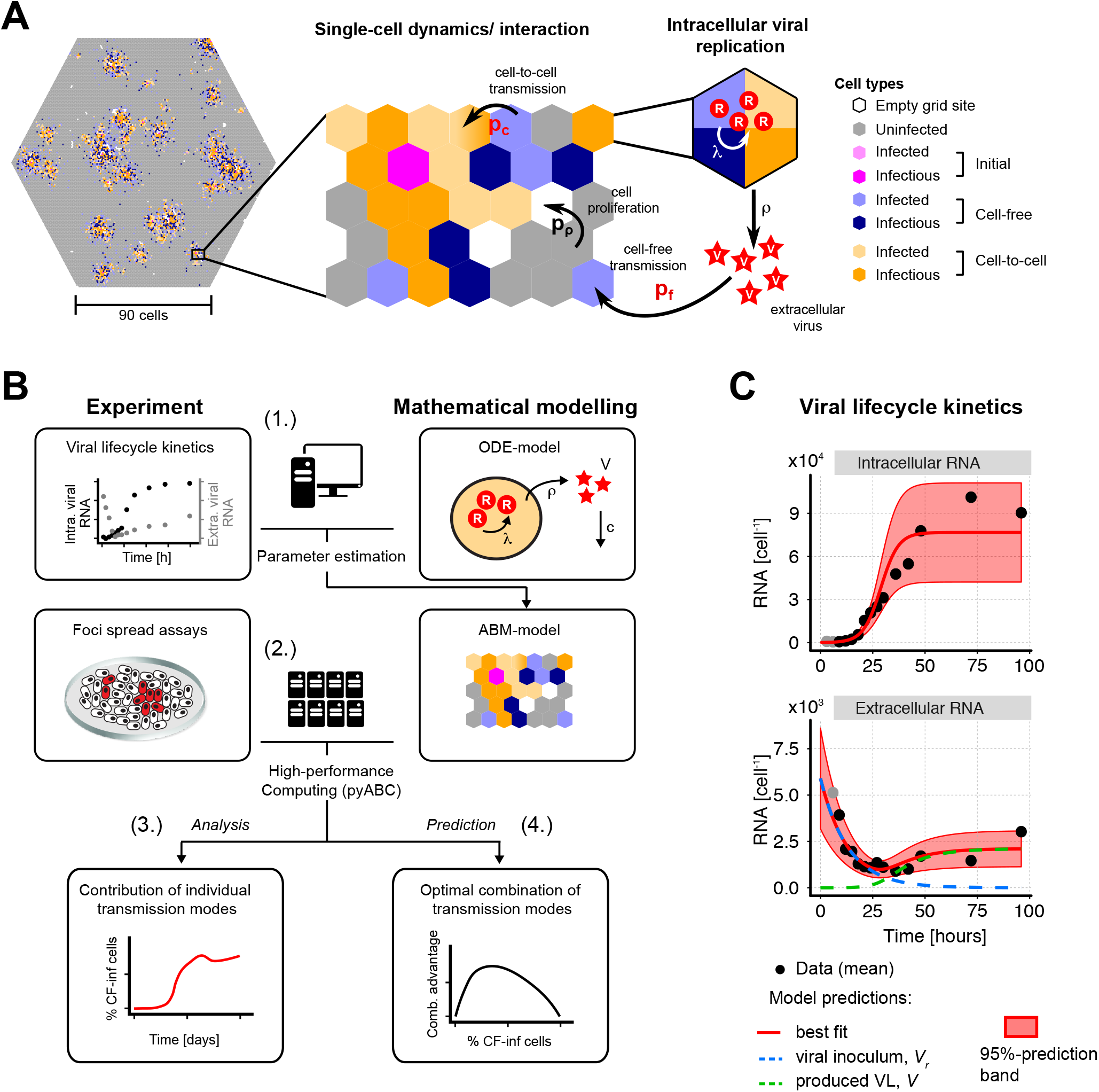
A multi-scale agent based model to describe HCVcc spread assay dynamics: (**A**) Schematic of the agent based model showing the multi-level structure considering single-cell and intracellular viral replication dynamics when simulating HCV spread. Individual cells are distributed on a hexagonal grid (left), which represents parts of the *in vitro* culture system. These uninfected cells are able to proliferate to fill empty grid sites with probability *p_p_*. Starting from a number of initially infected cells, uninfected cells can get infected by cell-free or cell-to-cell transmission by stochastic processes according to the probabilities *p_f_* and *p_c_*, that depend on the extracellular viral concentration, *V*, or the intracellular viral load, *R*, of infectious cells, respectively. Intracellular viral replication within these cells considers the changing concentration of intracellular viral RNA, *R*, and viral production, ρ (right). See *Materials & Methods* for a detailed description of the different processes considered. (**B**) Sketch of the stepwise analytical approach used to infer the contribution of the different transmission modes to the HCV infection dynamics: (1.) Experimental data on viral lifecycle kinetics are combined with a mathematical model to quantify and parameterize intracellular viral replication and viral export. (2.) These results are then incorporated in the multi-scale agent based model (ABM) that is used to simulate HCV spread dynamics under the various experimental conditions. The ABM is fitted to the time-resolved focus size distribution data with and without the use of anti-E2 using a high-performance computing approach (pyABC). (3.) The parameterization of the individual processes within the ABM to describe the observed dynamics allows us to infer the contribution of the individual transmission modes to viral spread given different scenarios. (4.) In addition, based on the obtained simulation environment, we predict the advantage of combined transmission modes for viral spread. (**C**) Experimental data (black dots) and model predictions (red line) showing the dynamics of intra- and extracellular viral RNA over time. Red-shaded area indicates the 95%-prediction interval of model predictions. For the extracellular RNA (lower panel), the dashed lines indicate the contribution of the initially applied (blue) and newly produced (green) virions to the total viral load.

To apply the multi-scale mathematical model to our experimental data, we used a step-wise approach (**Fig. 3B**). In the first step, we parameterized the intracellular viral replication dynamics by fitting the corresponding deterministic mathematical model to the life cycle kinetics data from **Figure 1**. Parallel cultures from this experiment were stained at 30 and 72 hours p.i. to determine the percentage of cells infected and other cultures were utilized to count cells per well at 0, 36, 72 and 96 hours p.i.. which allowed us to estimate the average HCV RNA copies per individual infected cell. The model provides a good description of the observed infection kinetics (**Fig. 3C**) with the maximal production rate of intracellular RNA estimated at λ = 0.24 [0.194, 0.278] h^-1^, the concentration of viral RNA reaching a carrying capacity of *R_C_*=7.73 [6.30, 9.76] x 10^4^ RNA copies per cell, and a viral export of ρ=2.10 [1.24, 3.47] x10^-3^ h^-1^ (numbers describe best estimate and 95%-confidence intervals of estimates) (see also **Table S1**). This parameterization of the intracellular replication dynamics within individual cells is then used in the second step, in which we applied the whole multi-scale model to the HCV spread assay data in order to parameterize the kinetics of cell-free and cell-to-cell transmission, the viral diffusion rate, and the loss of anti-E2 neutralization efficacy in the treated cultures. In this second step, we used a distributed, likelihood-free simulation-based method based on approximate Bayesian computation (pyABC,[18], see *Materials and Methods*) to fit the stochastic, computationally demanding models to the experimental measurements. Before analyzing the actual experimental data, we validated the general appropriateness of our approach by simulating data in correspondence to the experimental measurements and testing the ability of our method to retrieve the parameters used for simulation. Our analysis showed a correct recovery of the predefined parameters using focus size distribution of simulated treated and untreated HCV spread assays for model adaptation (**Fig. S2**).

### Mathematical analysis allows determination of transmission kinetics and reveals varying contributions of viral transmission modes to HCV spread

After validation of the general applicability of our approach, we fit our agent-based model to the HCV spread data, separately analyzing the two sets of experiments represented in **Figure 2B** (Exp. A) and **2C** (Exp. B). Corresponding to the experimental scenarios, model simulations were run with or without simulating the neutralization of cell-free HCV via anti-E2. To account for the stochasticity due to the different number of wells within the individual experiments, we used the mean of two (Exp. A) and five (Exp. B) individual ABM-simulations for comparison against the experimental data. Each run by pyABC evaluated ~21,000 particles, i.e., parameter combinations, with each particle comprising the corresponding number of individual simulations. The algorithm was stopped after 13-15 generations, i.e., successive improvements of the approximation of the parameter posterior distribution (see *Materials and Methods*) as sufficient convergence was reached. With these methods, our multi-scale model is able to reproduce the observed experimental data. Specifically, the model recapitulates the focus size distribution in both untreated (**Fig. 4A** and **Fig. S3A**) and anti-E2 treated cultures (**Fig. 4B** and **Fig. S3B**), except for a tendency to predict a higher number of small foci sizes particularly at later time points. Total infected cell numbers are generally well predicted for both conditions and experiments (**Fig. 4C-D** and **Fig. S3C-D**). Thus, our agent-based model is able to reproduce the experiments and mimic the complex spatio-temporal dynamics on a single-cell level. As would be expected, model estimates for the coupling parameter for the change of viral concentrations between grid sites were similar between the two experiments and in the order of 10^-2^, which corresponds to effective viral diffusion coefficients, *D*, of 10^-2^ to 10^-1^ μm^2^/s (**Fig. 4E** and **Table 1**). However, our analysis predicted a higher usage rate of anti-E2 within the later experiment (Exp. B) compared to the first round of experiments (Exp. A), which is consistent with the hypothesis that there was considerable anti-E2 escape in Exp B. Estimates for the model parameters determining the infectivity of HCV intracellular viral material indicated that the time-span between two successful cell-to-cell transmission events originating from the same infected cell was in between 23 and 42 hours. This intracellular viral infectivity in combination with high intracellular RNA levels impairs the identifiability of the cell-to-cell transmission factor β_c_ that scales the probability of infection with the amount of intracellular viral RNA for each time-step and corresponds to the cell-to-cell transmission rate (**Fig. S4** and **Fig. S5**). As such, the parameter combinations to describe the experimental data for both experiments contained estimates for β_c_ that spanned a broad range between ~10^-6^ – 10^-2^ min^-1^ intraRNA^-1^. In contrast, parameter estimates for the cell-free transmission factor were in a tighter estimated range and varied between the two experiments with β_f_ ~ 10^-4.5^ – 10^-3^ min^-1^ extraRNA^-1^ in Exp. B compared to β_f_ ~ 10^-6.5^ – 10^-5^ min^-1^ extraRNA^-1^ in Exp. A (**Fig. 4E** and **Table 1**). Independent of the individual parameter estimates, all obtained parameter combinations provided a robust prediction of the contribution of the individual transmission modes to viral spread. For Exp. A, which had less cell division, the initial phase of viral spread up to 1-1.5 days p.i. is almost exclusively characterized by cell-to-cell transmission even in cultures that are not treated with anti-E2 (**Fig. 4F**). Cell-to-cell transmission remained the dominant mode of spread with on average ~76% [63%, 89%] of successful infections mediated by this transmission mode at 96 hours p.i. (Numbers in brackets indicate min and max. predictions by selected parameter combinations, see **Fig. 4F**). Cell-free transmission is predicted to also contribute to viral spread in the anti-E2 treated cultures starting around ~2 days p.i. and being responsible for ~9 % [0%, 25%] of all infections 96 hours p.i.. Regardless, cell-to-cell transmission is predicted to be the dominant mode of transmission in Exp. A. In contrast, for Exp. B, which cell division was more significant, the contribution of the individual transmission modes is predicted to change over the time course of the experiment, with most of the infections (~72% [62%, 80%] (untreated) and ~72% [64%, 81%] (anti-E2 treated) being mediated by cell-free transmission at later time points (**Fig. 4G**). Thus, the virus shows versatility in the contribution of the individual transmission modes to the progression of infection between the experimental conditions.

**Figure 4:**
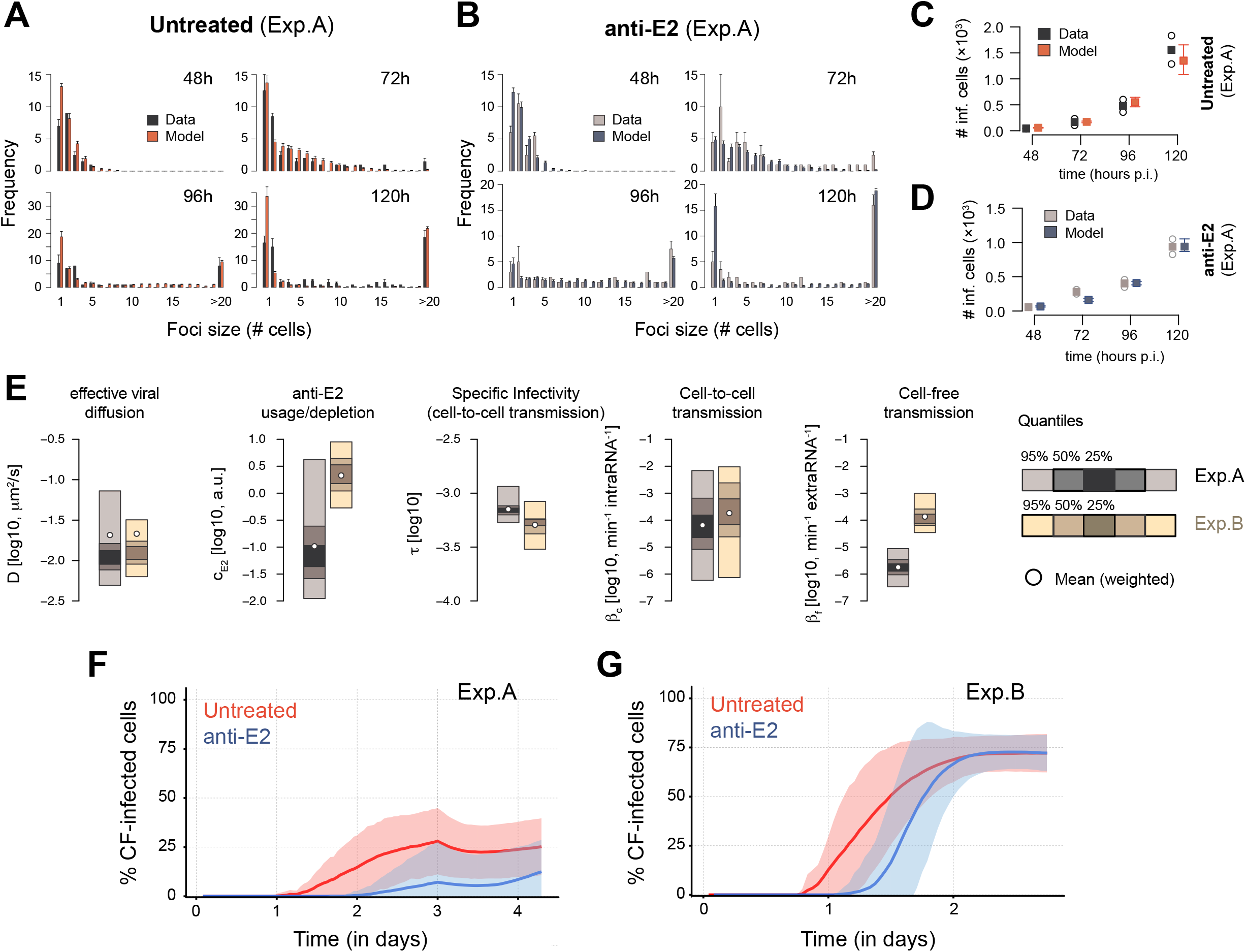
Model fits, parameter estimates and relative contribution of cell-free and cell-to-cell transmission modes to HCV spread: (**A,B**) Measured (black/grey) and predicted (red/blue) focus size distributions for Exp. A in the absence (**A)** and presence (**B**) of anti-E2 at 48, 72, 96 and 120 hours p.i. and calculated across 2 replicates (=wells) according to the experimental conditions. Predictions are based on an exemplary parameterization of the model as obtained by the fitting procedure. **(C,D)** Measured (black/grey) and predicted (red/blue) average number of infected cells in untreated (**C**) and anti-E2 treated (**D**) wells calculated across 2 replicates (=wells). (**E**) Credibility intervals for the individual parameter estimates obtained by the high-performance computing approach (pyABC) for Exp. A (black/grey) and Exp. B (orange/wheat) after 15 generations of optimization. White circles indicate the weighted mean for each parameter. Corresponding estimates are shown in Table 1. **(F,G)** Predicted proportion of cells infected by cell-free transmission for Exp. A (**F**) and Exp. B (**G**) over the course of the experiment. Mean (solid lines) and 95%-CI (shaded area) as calculated from all model predictions obtained from the best performing parameter combinations (=particles) with a distance smaller than 4.0, i.e. 10-15 particles, for untreated (red) and anti-E2 treated (blue) simulations.

**Table 1:** Parameter estimates for the different experiments. The estimate for each experiment indicates the weighted mean out of all parameter combinations obtained after 15 (Exp. A) and 13 (Exp. B) generations, i.e., optimization repeats, of pyABC (see *Materials and Methods)*. Numbers in brackets indicate 95% credibility intervals.

| Description | Parameter | Unit | Experiment |  |
| --- | --- | --- | --- | --- |
|  |  |  | Exp. A | Exp. B |
| Cell-to-Cell transmission rate (i.e., scaling factor) | $\beta_c$ | $\log_{10}, \text{min}^{-1} \text{intraRNA}^{-1}$ | -4.19<br>[-6.24, -2.15] | -3.74<br>[-6.13, -2.02] |
| Cell-free transmission rate (i.e., scaling factor) | $\beta_f$ | $\log_{10}, \text{min}^{-1} \text{extraRNA}^{-1}$ | -5.75<br>[-6.48, -5.05] | -3.87<br>[-4.46, -3.00] |
| Rate of anti-E2 usage/depletion due to neutralization of extracellular RNA | $c_{E2}$ | $\log_{10}, \text{arbitrary unit min}^{-1}$ | -0.98<br>[-1.95, 0.62] | 0.33<br>[-0.27, 0.95] |
| Specific infectivity for cell-to-cell transmission | $\tau$ | $\log_{10}$ | -3.15<br>[-3.27, -2.93] | -3.29<br>[-3.52, -3.07] |
| Viral diffusion coefficient* | D | $\log_{10}, \mu\text{m}^2/\text{s}$ | -1.68<br>[-1.96, -0.91] | -1.67<br>[-1.91, -1.27] |

\* The theoretically derived diffusion coefficient D according to the Stokes-Einstein equation is $D=8.05 \mu\text{m}^2/\text{s}$ ( $= 0.91$ on a $\log_{10}$ scale) according to the assumed diameter of a hepatocyte ( $20 \mu\text{m}$ ), of a HCV virion ( $30 \text{ nm}$ ) and the dynamic viscosity of the medium. The estimated diffusion is much slower than the theoretical prediction as it represents the effective diffusion allowing exported virions to reach uninfected cells in the culture.

### Simultaneous occurrence of cell-free and cell-to-cell transmission enhances viral spread

As inferred from the experimental data and shown by our mathematical analyses, the spread of HCV relies on the simultaneous occurrence of both transmission modes. This suggests that the combined occurrence of cell-free and cell-to-cell transmission provides an advantage for viral spread and the progression of infection. Based solely on the number of cells that became infected in the experiment Exp. A in the absence and presence of neutralizing anti-E2, we calculate that the presence of cell-free spread resulted in a ~1.7-fold higher average number of infected cells/well at 120 hours p.i. (1559.5 (untreated) vs. 937.5 (anti-E2)). Using our multi-scale agent based model with the obtained parameterizations for this experiment to predict the infection dynamics given each transmission mode separately, we would only expect a 1.1-fold increase in the total number of infected cells with an average of 22% of cells being infected through cell-free transmission (**Table 2**). Thus, the combined occurrence of both transmission modes provides a substantial synergistic effect for viral spread that exceeds the simple additive contribution of both transmission modes. In order to determine the degree to which different ratios of cell-free and cell-to-cell transmission affect this advantage, we tested the effect of varying transmission modes in our multi-scale model. To this end, we simulated viral spread using either a “wild-type”, WT, strain that is able to spread by both cell-free and cell-to-cell transmission or two “mutant strains”, MUT-CF and MUT-CC, that are only able to spread by cell-free or cell-to-cell transmission, respectively. Comparing the spread dynamics of these three hypothetical viruses allowed us to assess the synergy achieved when varying ratios of cell-free and cell-to-cell transmission occur simultaneously. Hereby, we defined a relative synergistic effect, *RSE*, determined by the ratio of the number of infected cells obtained in each scenario, i.e., *RSE*=*I*_WT_/(*I*_WT_ +(*I*_MUT-CF_ + *I*_MUT-CC_)). An *RSE* of 0.5 would mean no synergistic effect of the simultaneous occurrence of both transmission modes, while values close to 1 indicate a large synergistic effect (see also *Materials and Methods*). We tested various scenarios for the combined probability of both transmission modes, also assuming different rates of viral diffusion (**Fig. 5A**). We find that for comparable viral diffusion rates as in our experiments (**Fig. 4E**), the relative synergistic effects are largest if ~60-70% of the infections are due to cell-to-cell transmission (**Fig. 5B, Fig. S6**), comparable to the cell-to-cell spread contribution predicted for Exp. A (**Fig. 4F**). Thus, under the experimental conditions with low cell division, HCV spread seems to use the optimal combination of both transmission modes for viral spread.

**Table 2:** Advantage of combined spread: Mean number of infected cells for experimental (Exp. A) and simulated data with and without anti-E2 treatment 120 hours post infection. Experimental data allowing for both transmission modes show a 1.66-fold increase in infected cells compared to anti-E2 treated cultures. Similar results for simulated data using the parameterizations obtained for Exp. A, with ~22% of infected cells being due to cell-free transmission. Simulations show that the additive combination of infected cells by cell-free and cell-to-cell transmission would only lead to a ~10% increase in infected cell numbers compared to anti-E2 treated cultures. (*The results for the ABM show the average over 10 individual simulations.)

|  | Treatment | Infected cells<br>(mean) | Fraction of cells<br>infected by CF | Fold Increase | Expected Fold<br>Increase |
| --- | --- | --- | --- | --- | --- |
| **Experiment**<br>**(Exp.A)** | anti-E2 | 937.5 | - | 1 | - |
|  | untreated | 1559.5 | - | 1.66 | - |
| **Simulation\***<br>**(ABM)** | anti-E2 | 1028 | 0 | 1 | - |
|  | untreated | 1650 | 0.22 | 1.61 | 1.09 |

**Figure 5:**
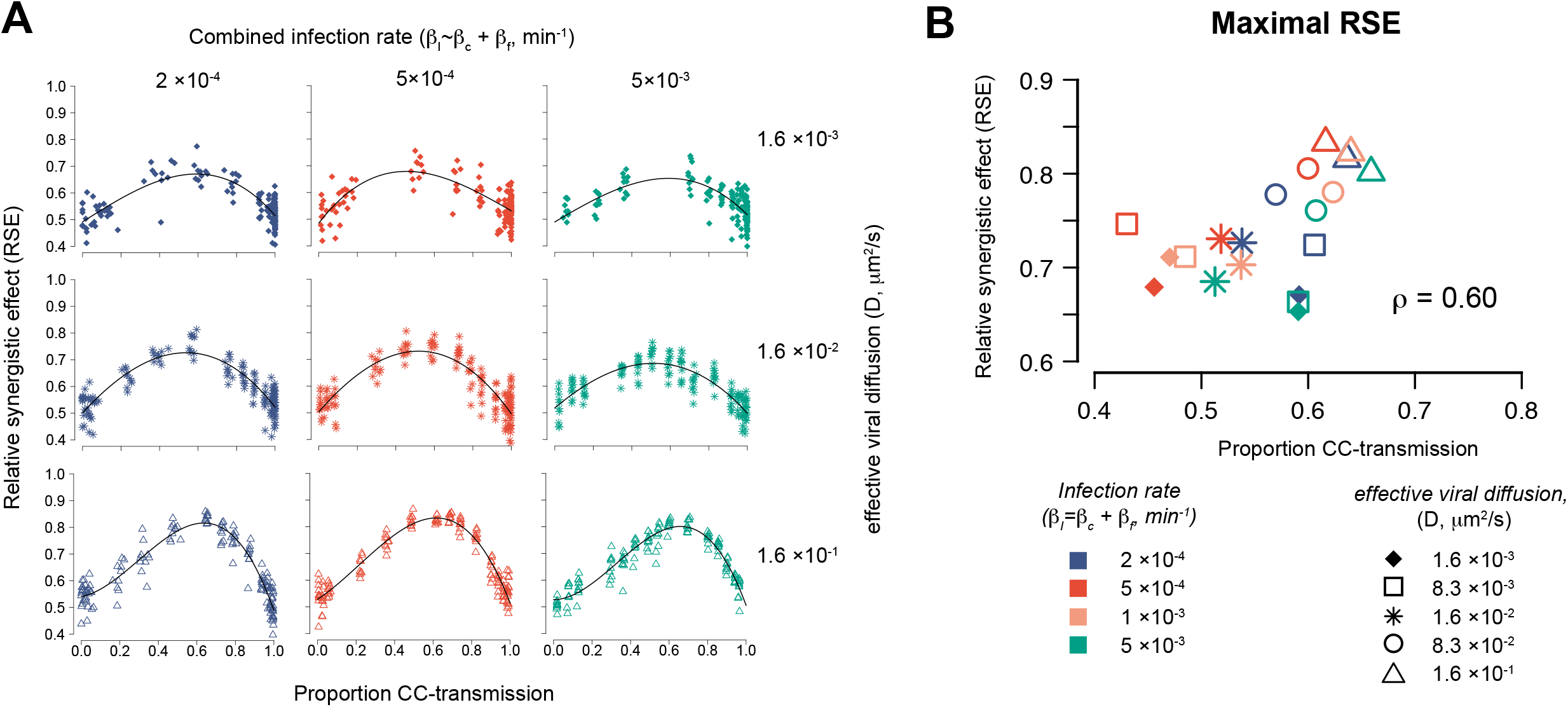
Advantages of combined modes of viral spread: **(A)** Predicted advantage of combined modes of viral spread indicated by the Relative Synergistic Effect *(RSE)* of combined viral spread. The *RSE* is shown dependent on the proportion of infections by cell-to-cell transmission given different viral diffusion coefficients and infection rates. Individual points indicate the maximal *RSE* obtained for simulating viral spread in the ABM for a 10 day time course with different combinations of the transmission factors for each transmission mode, β_*c*_ and *β_f_*, defining the combined infection rate *β_I_*. Curves show the result of a spline function of 3^rd^ degree fitted to the individual data. **(B)** Maximal *RSE* value and proportion of infections due to cell-to-cell transmission at which the fitted curves reached their maximum for different conditions analyzed (**A**, **Figure S6**)

## Discussion

Viral spread within a host is a critical parameter that determines the kinetics of infection and the efficacy of antiviral therapies. Cell-to-cell spread has specifically been implicated in the establishment of persistent infections [19], the propagation of antiviral resistant mutants [8], and in requiring increasing effective doses of antiviral drugs [19]. Yet, many aspects of viral cell-to-cell spread, and the relative contributions of cell-free versus cell-to-cell transmission during infection are still unknown. Here, we present a cross-disciplinary approach that combines experimental kinetic data and multi-scale mathematical modeling to determine HCV spread kinetics, and to disentangle the contribution and interplay of cell-free and cell-to-cell transmission modes.

While the experimental data demonstrated that infection progressed faster when both transmission modes were available for spread relative to when cell-free transmission was inhibited by anti-E2 (**Fig. 2D),** our mathematical analysis revealed additionally that the combined spread by cell-free and cell-to-cell transmission results in a synergistic effect that exceeded the additive contribution of both transmission modes (**Table 2**). Conceptually, synergy between the two modes of spread is expected as the fraction of cells contributing to cell-to-cell spread within individual foci is limited by whether or not infected cells are adjacent to uninfected neighboring cells. This fraction decreases as foci expand with more and more cells enclosed by other infected cells. As such, new foci formed by cell-free spread seed additional areas for cell-to-cell spread to occur, a concept that is also corroborated by the increased number of individual foci observed at later time points in our experiments (**Fig. 2**). In experiments with low cell division (represented by Exp. A), we also observed that largest foci were formed when both means of HCV spread were allowed to proceed, indicating that cell-free spread could also contribute to individual foci growth. However, this could be influenced by the *in vitro* cell culture system in which cells are maintained in relatively stagnant media. To which extent this contribution might also play a role *in vivo*, where blood flow influences viral dispersion remains to be determined.

Combining our experimental data with a mathematical model allowed us to quantify the contribution of cell-free and cell-to-cell transmission modes throughout the time course of the experiments. While the initial predominance of cell-to-cell spread observed in our earlier *in vitro* culture experiments (**Fig. 4F**) is consistent with the idea that cell-to-cell spread was the predominant mechanism of HCV transmission, after switching lots of FBS we observed an increase in the number of cell divisions occurring during our experiments, as well as an increase in the degree of cell-free spread (**Fig. 4G**). These results suggest that the contribution of each transmission mode may be influenced by the environmental conditions encountered. The observation that viral transmission modes are influenced by environmental conditions is in line with observations made for other viruses, such as HIV-1 [20, 21] and highlights the utility of trying to incorporating more physiologically relevant conditions in future experiments when studying viral spread [22].

Arguably, cell proliferation dynamics could have an impact on the efficacy of viral transmission modes. One could envision that actively dividing cells do not maintain the necessary stable cell-cell contacts required to mediate cell-to-cell virus transmission. If this is the reason for the inferred increase in cell-free spread in more actively dividing cell cultures (**Fig. 4G**), this would suggest that there might be a predominance of HCV cell-to-cell spread in the liver where hepatocyte division is generally low. However, the more actively dividing cell cultures also presumably had higher extracellular virus levels, as indicated by the notable anti-E2 escape in our *in vitro* data and confirmed by our mathematical analyses that estimated higher rates of anti-E2 usage in Exp. B compared to Exp. A (**Table 1** and **Fig. 4E**). Thus, there could also be a self-reinforcing effect on the importance of cell-free transmission, with small cell-free infection advantages in high proliferating cell cultures leading to higher extracellular viral levels, and, thus, enhanced contribution of cell-free transmission to viral spread. To determine if and how the observed differences in cell proliferation are responsible for the observed changes in ratio between the different modes of spread, additional experimental analyses will be needed.

Besides influencing cell proliferation dynamics, further variables that might differ between the two serum lots could affect the efficacy of individual transmission modes and might be relevant to environmental changes within the liver of chronically infected individuals (as e.g. influenced by diet, medication, age, gender, and liver health). For example, different concentration of lipids could affect HCV entry and replication dynamics [23–25]. In particular, because HCV is known to exploit the very low density lipoprotein pathway and lipid droplets for viral assembly and maturation [26], lipid-enriched environments could potentially enhance cell-free transmission.

Independent of environmental conditions, the predicted stability of the inferred ratios of cell-free vs. cell-to-cell transmission at later time points despite the ongoing increase in the number of infected cells (**Fig. 4F-G**) indicates that these ratios seem to be favorable for viral spread in the respective experimental scenario. This raises the question whether there is an optimal balance of these transmission modes and whether our *in vitro* infections reflected that optimal ratio. Simulating viral spread for different ratios and efficacies of cell-free vs. cell-to-cell spread in conditions experiencing slow cell division, we found that the relative synergistic effects are largest if ~60-70% of infections are due to cell-to-cell transmission (**Fig. 5**). Notably, this is comparable to the transmission mode contributions estimated for Exp. A. This suggests that HCV naturally achieves the ratio of transmission modes that optimally exploits the available synergistic advantages even within the relatively non-competitive *in vitro* culture dish.

It is tempting to speculate that a dominant contribution of cell-to-cell transmission could also play a role in HCV spread *in vivo*. Indeed, detailed analyses of liver biopsy samples of patients chronically infected with HCV by single cell laser microdissection and microscopy techniques, revealed that infected cells occurred in clusters that are heterogeneously distributed throughout the tissue [27–29]. These patterns of infection support the simultaneous occurrence of both transmission modes *in vivo* [30]. Considering the various immune responses that HCV encounters *in vivo*, cell-to-cell transmission might be even more beneficial *in vivo* than what was measured here *in vitro*.

Our analysis combining experimental data and mathematical modeling allows us to quantitatively assess HCV transmission dynamics on a cellular level. However, while our mathematical model is able to provide a good representation of the observed spread dynamics, it generally overestimates the number of very small foci sizes (i.e., single cell foci) at later time points (**Fig. 4** and **Fig. S3**). There are several possible explanations for this discrepancy. One possible explanation is the limited ability of the grid-based model to completely account for experimental cell proliferation dynamics. *In vitro*, even when cells are plated at confluence, they continue to divide while decreasing in size as they pack in tighter within the allotted space. With simulated cells having a fixed volume, our model does not allow for continuous cell proliferation once the grid has been filled, limiting the ability to account for foci growth due to cell division. Advanced modeling frameworks, such as cellular Potts models that allow for dynamic cell shapes [31], as well as spread assay experiments within non-dividing cell cultures could help to address this discrepancy. Another theoretical possibility for the observed overestimation of small foci sizes is that defective particles initiate non-productive infections that result in non-expanding single-cell foci. This could cause the model to overestimate the contribution of cell-free transmission at earlier time points, leading to the increased predicted frequency of newly founded foci at later time points. Notably, in this case, the contribution of cell-free transmission to viral spread might be even smaller than estimated here.

Besides these limitations, the current model is quite complex, such that individual parameter estimates have to be taken with care. For example, estimates for the individual parameters describing viral transmission, anti-E2 usage and effective viral diffusion can vary over different orders of magnitude. However, predictions regarding the contribution of the transmission modes to viral spread by the determined parameter combinations are quite robust, allowing for a reliable assessment of these quantities. All parameters governing the viral kinetics in the agent-based model were identifiable and the best fit estimates were adopted for the purpose of describing HCV spread in a monolayer of Huh7 cells (**Table S1**). We used a simplified model system that explains HCV viral replication and export dynamics in a continuous and deterministic way, providing an appropriate representation of the observed lifecycle dynamics and the development of intracellular and extracellular viral loads (Figure 3C). Although the initial intracellular RNA concentration was considerably larger than 1, presumably due to co-infection, abortive infections, and a significant association of non-infectious particles on the outside of the cells, these results do not bias the conclusions drawn from the agent-based model as the total concentration of intra- and extracellular viral RNA was always scaled with the transmission factors, *β_c_* and *β_f_*, respectively, which were estimated using the observed focus size distributions.

Additional experimental advancements, such as automated image analyses could increase the amount of available data and, thus, improve parameter inference in our mathematical models. In addition, experimental approaches that exclusively block cell-to-cell transmission would be desirable. However, both transmission modes rely on many of the same cell surface receptors [9, 11, 14, 32], making this currently experimentally difficult.

In summary, we monitored HCV spread kinetics *in vitro* and combined the experimental data with a multi-scale mathematical model to disentangle the contribution and interplay of cell-free and cell-to-cell transmission modes during viral spread. Our analysis revealed varying contributions of transmission modes to HCV spread under different culture conditions highlighting the adaptability of the virus. Regardless of environmental effects, our analysis also revealed synergistic effects between the two modes of transmission that seem to be optimally exploited during viral spread. This leads to the possibility that some of the advantages typically attributed specifically to cell-to-cell spread (i.e., the ability to establish viral persistence) may be due to having two synergistic modes of transmission rather than cell-to-cell spread itself.

## Materials and Methods

### Experimental Methods

#### Cells and Virus

Huh7 human hepatoma cells were obtained from F.V. Chisari (The Scripps Research Institute, CA) [33] and cultured in complete Dulbecco’s modified Eagle’s medium (cDMEM) supplemented with 100units/ml penicillin, 100mg/ml streptomycin, 2mM L-glutamine (Corning), 10mM HEPES (Santa Cruz), and non-essential amino acids (Thermofisher Scientific) and 10% fetal bovine serum (FBS) (Hyclone or Gibco). Cells were maintained at 37°C in 5% CO_2_. Stocks of HCV were produced from a plasmid encoding the JFH-1 genome that was provided by Takaji Wakita (National Institute of Infectious Diseases, Tokyo, Japan) [33, 34]. Methods for HCV RNA *in vitro* transcription and electroporation into Huh7 cells have been described previously [35]. Media collected from HCV RNA transfected cells was then used to infect Huh7 cells at a multiplicity of infection (MOI) of 0.01 foci forming units (FFU)/cell. Culture media from those infections were harvested, pooled, aliquoted, frozen, and titered as described previously [35]. To achieve high titer virus stocks for high MOI experiments, virus was collected in serum-free, phenol red-free media and concentrated via Amicon ultra centrifugation filters (Milipore) prior to aliquoting and freezing.

#### Reagents

The human anti-HCV E2 antibody (AR3A) was provided by Mansun Law (The Scripps Research Institute, CA)[36]. Mouse anti-HCV NS5A (9E10) was provided by Charlie Rice (Rockefeller University, NY)[37]. Secondary antibodies anti-human-HRP and goat anti-mouse-HRP were purchased from Thermofisher Scientific and Vector Labs, respectively. The 3-amino-9-ethycabazole (AEC) substrate kit was purchased from BD Pharmingen.

#### High MOI HCV Life cycle Kinetics

Huh7 cells were plated in 96 well plates at 4,000 cells per well. The next day cells were inoculated with serum free HCV at a MOI of 6 in 50μL of serum-free cDMEM for 3 hours. The inoculum was then removed, and the wells were rinsed twice with warm cDMEM before adding 200μL fresh cDMEM with 10% FBS. Media and cell lysates were collected from triplicate wells at 0, 3, 6, 9, 12, 15, 18, 21, 24, 27, 30, 36, 42, 48, 72 and 96 hours post inoculation (p.i.). To determine the amount of cell division, cells in parallel wells were counted at 0, 36, 72 and 96 hours post-inoculation. Additionally, duplicate wells were fixed at 30 and 72 hours p.i. to determine the percent infected cells by immunostaining for HCV.

#### RNA Isolation and Quantification

Total intracellular RNA was isolated using an ABI PRISM 6100 Nucleic Acid Prepstation (Applied Biosystems), using the manufacturer’s instructions. Extracellular RNA was extracted from culture media spiked with 1μg mouse liver RNA which served as an internal extraction efficiency control RNA, using a KingFisher Duo Prime Purification System or a KingFisher Flex purification system (ThermoFisher), per the manufacturer’s instructions. Isolated RNA was used to create cDNA via random prime reverse transcription (Revertaid transcriptase, Thermofisher). Quantitative PCR was then performed with iTaq Universal SYBR Green Supermix (BioRad) using Applied Biosystems 7300 real-time thermocyclers. The thermal cycling program included an initial 30 second 95°C denaturation step followed by 40 cycles of denaturation (15 seconds at 95°C) and 1-minute annealing/extension step at 60°C. HCV copies were quantified relative to a serially diluted standard curve of the pJFH-1 plasmid. Intracellular HCV copies were normalized to cellular huGAPDH and extracellular HCV copies were normalized to mGAPDH. The PCR primers used to amplify HCV were 5′-GCC TAG CCA TGG CGT TAG TA −3′ (sense) and 5′-CTC CCG GGG CACTCG CAA GC-3′ (antisense). The PCR primers used to amplify GAPDH were 5′-GAA GGT GAA GGT CGG AGT C-3′ (sense) and 5′-GAA GAT GGT GAT GGG ATT TC-3′ (antisense).

#### HCV Titer Assay

Huh7 cells were plated at 4,000 cells per well in a 96-well plate. Approximately 24 hours later, 100μL of 10-fold serial dilutions of virus samples were added to the cells in duplicate. At 24 hours p.i., a 0.5% methylcellulose overlay was added to the wells. At 72 hours p.i., cells were fixed with 4% paraformaldehyde for 20 minutes, washed with 1X phosphate buffered saline (PBS) and immunostained for HCV. Titers (FFU/mL) were determined by counting the number of foci in multiple dilutions.

#### HCV Immunohistochemical Staining

Fixed cells were permeabilized by adding a 1:1 mixture of −20°C methanol:acetone for 10 minutes. After a 1X PBS wash, endogenous peroxidases were inactivated with 0.3% (v/v) hydrogen peroxide for 5 mins followed by another 1xPBS wash. Cells were blocked for 60 mins at room temperature on an orbital rocker with blocking buffer [1X PBS containing 0.5% (v/v) TritonX-100, 3% (w/v) bovine serum albumin (BSA) and 10% (v/v) FBS], followed by incubation with the primary antibody diluted in binding buffer [1X PBS containing 0.5% (v/v) TritonX-100 and 3% w/v BSA] for 60 mins at room temperature. Cells were incubated with mouse anti-HCV NS5A (9E10) (1:500) or human anti-HCV E2 (AR3A) (1:750), as indicated. After washing with 1X PBS, appropriate secondary antibody in binding buffer was added for 60 mins at room temperature. Secondary antibodies included either goat antimouse HRP (1:4) or goat anti-human HRP (1:750). After washing with 1X PBS, HRP staining was developed using an AEC substrate kit. The wells were washed with ddH2O and 1:1 glycerol:water mixture was added to the wells for storage.

#### Spread Assay

Details of the protocol have been previously published [9, 10] but briefly confluent monolayers of Huh7 cells in 96-well plates were infected with indicated FFU per well. After 17 hours incubation at 37°C, the inoculum was removed and media containing 1% dimethyl sulfoxide (DMSO) was added to slow cell growth because it has been previously shown that culturing Huh7 cells in 1% DMSO causes cell proliferation to stop after approximately 6 days [35, 38]. Cells were either left untreated or treated with neutralizing HCV E2 antibody (AR3A) at 10μg/ml, which has been documented to block HCV cell-free spread under the experimental conditions utilized herein [4, 9, 10]. Media was changed at 72 hours p.i., and every 24 hrs thereafter unless noted otherwise. The number of cells per well was counted at each time point throughout the course of the assay to assess cell division. Cells were fixed at 48, 72, 96 and 120 hours (Exp. A) or 48, 59, 72 and 83 hours (Exp. B) as indicated using 4% paraformaldehyde and immunohistochemically stained for HCV (with either anti-NS5A or anti-E2). The number of foci and foci sizes (i.e. cells/focus) were counted using light cell microscopy.

#### Quantifying Foci Size

Initially, the number of cells/focus was determined either by manual counting through a light microscope or by taking pictures with a Nikon Diaphot TMD inverted phase contrast microscope equipped with an Olympus DP21 camera and subsequently using Microsoft Paint Program to count the number of cells per focus in pictures (**Fig. 2A**). We expedited our quantification procedure by using the measurement tool in ImageJ to measure the area of representative cells at each time point (to account for decreasing cell size over time) as well as the area of each focus. For each time point, foci area was divided by the average area of single cells. As such, the majority of the foci size data was obtained via ImageJ quantification, which we confirmed to match the manual counting data (**Fig. 2A**).

### Mathematical modeling

#### Modeling viral life cycle kinetics

We described the intracellular and extracellular viral RNA kinetics for individual infected cells by modeling the dynamics of the corresponding concentrations. Intracellular HCV RNA concentration, *R*, is assumed to follow a logistic growth with a maximal replication rate λ and a total carrying capacity R_C_ for individual cells. Intracellular viral RNA is then exported at a constant rate ρ, to become new extracellular viral RNA, *V*. Furthermore, extracellular viral concentration is assumed to be lost at a rate c. The model is then described by the following system of ordinary differential equations:

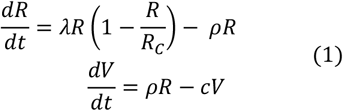

To allow parameter identifiability given the available data, the degradation of intracellular RNA was not explicitly modeled. Therefore, the parameter λ□ describes a net-replication rate considering viral production and degradation. The model in Eq. (1) was fit to the measured intracellular and extracellular RNA life cycle kinetics data (**Fig. 1A,B**) using a maximum likelihood approach. To account for the experimental setting, we additionally considered the initial viral inoculum, *V_r_*, that loses its infectivity at rate *c*, i.e., *dV_r_*/*dt*=-*cV_r_*. Thus, the measured extracellular viral RNA concentration, 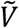, is a combination of *V_r_* and the newly produced viral RNA, *V, i.e*. 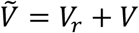. Despite removal of media, some viral particles adhere to hepatocytes resulting in high initial cell-associated RNA counts and residual viral particles in the media at early time points due to continuous binding and release. Therefore, measurements at 3 and 6 h p.i. were neglected in the fitting procedure. The 95-% confidence intervals of estimates were determined using the profile likelihood approach [39].

#### A multi-scale model to describe HCV infection dynamics

To analyze the HCV spread assays and determine the contribution of individual transmission modes, we developed a multi-scale model that describes HCV infection dynamics in a monolayer of cells. To this end, we extended a model that we had developed previously to simulate the spread of HCV in *in vitro* cell culture systems [14, 17]. The model accounts for the spatial distribution of cells and viral spread dynamics on a cellular level, as well as intracellular viral replication on a per cell basis and extracellular viral diffusion. Cells are placed on a two-dimensional lattice using a hexagonal grid structure, i.e., each cell having a maximum of six possible neighbors. For the whole grid, we assume closed boundary conditions with our simulation environment representing a single well. Extending the previously published model to additionally account for some degree of cell proliferation, a fraction of grid sites is left unoccupied in the beginning and uninfected cells are allowed to proliferate into empty adjacent grid sites following a normal distribution with mean μ and standard deviation σ=0.1. This fraction was set to 60% of to allow for cell division within the model, while also ensuring sufficient cell confluence and densely packed grids at later time points. For our simulations, we took a simplified approach for modeling cell division by averaging the cell division experimentally observed over the course of the entire experiment so that individual cells were assumed to divide every 32h (Exp. A) or 24 h (Exp. B) on average which was simulated stochastically for each cell. Cells are stationary and are infected by either cell-free or direct cell-to-cell transmission with the probability of infection depending on the concentration of cell-free virus at the respective grid site or the intracellular viral concentration of neighboring infected cells, respectively. Upon infection, intracellular viral replication and export follows the dynamics as described in Eq. (1) with the corresponding ordinary differential equations discretized and described by a set of difference equations with a time-step size of Δt=1 minute. Thus, Eq.(1) becomes to

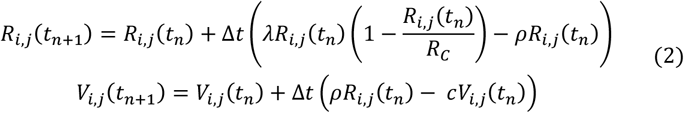

where *R_i,j_*(*t_n_*) and *V_i,j_*(*t_n_*) denote the concentration of intracellular and extracellular viral RNA for the cell at grid site (i,j) at time step *t_n_*. Exported viral RNA will contribute to the extracellular viral concentration at the respective grid site, *V_i,j_*. Diffusion of viral particles between grid sites follows the approach introduced by Funk et al. [40] with:

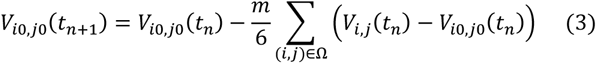

Hereby, *V_i0j0_(t_n_*+*1)* denotes the viral concentration at grid site (*i_0_,j_0_*) at time step *t_*n*+*1*_*, with *m* and Ω denoting the fraction of viral particles allowed to diffuse and the set of neighboring grid sites of (*i_0_,j_0_*), respectively. In the beginning of a simulation and following the experimental protocol, infected cells were initialized according to a truncated exponential distribution as described in [17].

##### Infection by cell-free and cell-to-cell transmission

The probability of a hepatocyte at position *(i,j)* to get infected by cell-free transmission in time-step *t_n_*, 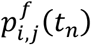, depended on the concentration of extracellular virus at the corresponding grid site, *V_i,j_(t_n_)*, and a scaling factor *β_f_* that corresponds to the cell-free transmission rate as used in deterministic mathematical models to describe viral spread [17, 20, 41]. Thus, at each time-step the probability for cell *(i,j)* to get infected was calculated by

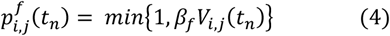

and a Bernoulli-trial with this probability was performed. In case of a successful infection, the extracellular viral concentration at this grid-site was reduced by R_0_. In case *V_i,j_(t_n_)*<R_0_, additional neighboring grid sites were considered to reduce the local viral concentration by R0.

Analogously, the probability of an infected cell to infect a neighboring uninfected cell by direct cell-to-cell transmission was calculated by

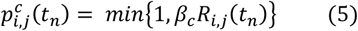

where *R_i,j_(t_n_)* denotes the concentration of intracellular viral RNA in cell *(i,j)* and *β_c_* the corresponding scaling factor representing the cell-to-cell transmission rate. If there was at least one uninfected cell in the direct neighborhood, a Bernoulli-trial with probability 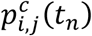 was performed. In case of a successful infection, the intracellular viral concentration in the infecting cell was reduced by R0. In addition, to account for possible unsuccessful cell-to-cell transmissions events despite a high intracellular RNA concentration due to non-infectious viral material (i.e. low specific infectivity), a factor τ was introduced, that delayed the occurrence of another transmission event originating from the same infected cell. This factor of intracellular HCV specific infectivity corresponds to the waiting time between two successful cell-to-cell transmission events from one infected cell and was therefore sampled from an exponential distribution with average τ. The delay was also considered for any newly infected cells before they are able to contribute to cell-to-cell spread.

##### Modeling anti-E2 treatment effects

As an additional extension to our previous model [17], we also added the concentration of anti-E2 as included within some of the experimental cultures. To this end, extracellular virus was reduced through neutralization by anti-E2 dependent on its relative concentration, *E2*, by

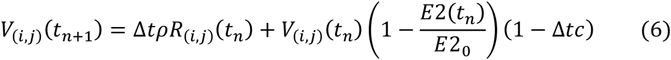

with *E2_0_* denoting the initial concentration of anti-E2 and *E2*(*t_n_*) the concentration at time point *t_n_*. Through the neutralization of virus, anti-E2 is depleted/consumed at rate *c_E2_*, reducing the concentration according to

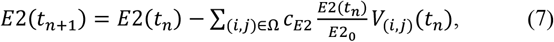

where Ω is the set of all grid sites. Note that we assumed a homogenous concentration of antiE2 throughout the grid. For simplicity, *E2* is given in arbitrary units with *E2_0_*=1.

The complete model was implemented in the C++ programming language. Simulations were performed using a time step, Δ*t*, of one minute. A sketch of the multi-scale model with its different elements is shown in **Fig. 3A.**

#### Parameter inference

The multi-scale model was fit to the experimental data to determine the rates for cell-free and cell-to-cell transmission, the intracellular HCV specific infectivity parameter τ, the rate of clearance of antiE2, *c_E2_*, and the effective viral diffusion, *D*, within these cultures. For model fitting, a likelihood-free simulation-based Approximate Bayesian Computation - Sequential Monte Carlo (ABC-SMC,[42]) method was employed, using sequential importance sampling to obtain an increasingly better approximation of the Bayesian parameter posterior distribution. Likelihood evaluation is circumvented by assessing the distance of the observed data to model simulations for sampled parameters, according to a distance measure (defined below), and accepting only particles below a sequentially reduced acceptance threshold. Fitting was performed to untreated and treated cultures simultaneously with all empirical spread data shifted by 18 hours, which was used as a lower bound to account for the delay between the time point of infection and experimental detection of infection. To account for the stochasticity due to the different number of wells used within the experiments, we used the mean of two (Exp. A) and five (Exp. B) individual simulations for comparison against the experimental data.

##### Distance measure used to fit the data

To capture the changing focus size distributions as well as the number of infected cells over the course of the experiment, we defined a relative distance between the predicted, *I_pred_*, and measured, *I_exp_*, total number of infected cells as follows:

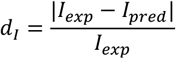

To describe the focus size distribution, the predicted, *f_pred_*, and measured, *f_exp_*, cumulative density functions for the occurrence of a focus with a specific size were calculated. Subsequently, the enclosed area between predicted and experimental cumulative density functions was divided by the average focus size as observed in the experiment

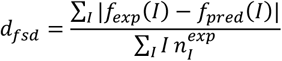

Here, the relative frequency of each focus size in the experiment is denoted by 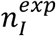. Dividing by the average focus size allowed comparable distances between early and late time points as the occurrence of large foci at late times would otherwise bias the calculated distance (**Fig. S6**). The sum of both distances, *d_total_* = *d_I_* + *d_fsd_* was then applied in the ABC-SMC algorithm. Due to the surprisingly fast increase of infected cells in antiE2 treated wells between 83 and 96 hpi in Exp. B in comparison to the slower increase in the untreated culture systems, the last time point was not considered within this analysis.

##### Parameter fitting by pyABC

To fit the agent-based model to the experimental data, we used the tool pyABC [18], which employs a distributed ABC-SMC algorithm based on [42]. The algorithm subsequently performs the following steps to find the best parameter set ν□(ν_□_,ν_□_,…,ν_□_) for explaining the data over a sequence of iterations *t*=*1*,…,*n_t_*:

1. Sample parameters from a proposal distribution 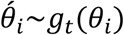
2. Simulate data from the model using the sampled parameter combinations, 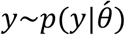 with 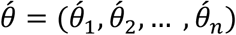.
3. Calculate the distance, *d*, between simulated and observed data and accept the parameter combination if the distance is below a given threshold, 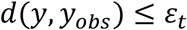.

Here, *g_1_* is the prior and subsequently *g_t_* is based on a multivariate normal kernel density estimate of the accepted particles (=parameter combinations) in the previous iteration, thus allowing to reduce the acceptance threshold *ε_t_* while maintaining high acceptance rates. The acceptance threshold was automatically chosen as *ε_t_* = median_i_(d_i_) of the accepted distances in the previous iteration, a strategy that has proven to be robust [43]. We used a population size of *N* = 100 or 200, meaning the algorithm needed to accept *N* parameter combinations according to the given threshold *ε_t_* before starting the next iteration. The procedure is automatized in the pyABC-framework [18] and customized for high performance computing systems. In particular, the framework uses dynamic scheduling to minimize the overall runtime on. Please refer to Klinger et al. [18] for more detailed information.

#### Evaluating the synergistic effect of simultaneous occurrence of cell-free and cell-to-cell transmission

In order to determine to which degree different ratios of cell-free and cell-to-cell transmission affect the spread synergy achieved by the combined occurrence of both transmission modes, we tested the effect of varying ratios of the transmission modes in our multi-scale agent based model. We simulated viral spread either using a “wild-type”, WT, strain that is able to spread by cell-free and cell-to-cell transmission or two “mutant strains”, MUT-CF and MUT-CC, that are only able to spread by cell-free or cell-to-cell transmission, respectively. We then defined the relative synergistic effect, *RSE*, of the simultaneous occurrence of both transmission modes by calculating the ratio of the number of infected cells obtained in each scenario, i.e., *RSE*=*I*_WT_/(*I*_WT_ +(*I*_MUT-CF_ + *I*_MUT-CC_)). The *RSE* is related to the expected fold-increase by *Fold*= (1/*RSE* – 1)^-1^. Thus, an *RSE* of 0.5 would mean no synergistic effect of the simultaneous occurrence of both transmission modes, while an *RSE* of 0.8 corresponds to a 4-fold higher number of infected cells in comparison to the separated spread of both mutants.

We simulated the dynamics for varying assumptions for the combined occurrence of cell-free and cell-to-cell transmission, *β_I_*=*βf*+*β_c_*, assuming 15 different combinations for the ratio of the transmission factors for cell-to-cell *β_c_*, vs. cell-fee infection, *β_f_* to cover different proportions of transmission modes. In addition, the impact of viral diffusion on the outcome was tested by varying the effective viral diffusion rate over different orders of magnitude. For each parameter combination, 10 simulations were performed simulating a time-period of 10 days. The equilibrium value of the proportion of cells infected by cell-to-cell transmission was determined for each simulation and plotted against the maximal *RSE* obtained. For better comparison, a 3^rd^ degree polynomial was then fitted against all 150 simulations for each combination (*pI,D*) (i.e. 15 ratios *β_c_*/*β_f_*, 10 simulations each) to determine the proportion of cell-to-cell transmission at which the *RSE* was maximal for the investigated condition. We considered 5 different values for *_pI_* and 4 different viral diffusion rate, *D*, in total (see **Fig. S5**).

## Statistical Analysis

Statistical comparison of foci sizes between untreated and anti-E2 treated cultures within the two experiments (Exp. A and Exp. B, **Fig. 2B,C**) were performed using Mann-Whitney U-test. Note that foci sizes from individual wells were combined for the analysis.

## Acknowledgments

We thank Elba Raimundez Alvarez for expert technical help with pyABC. For high-performance computational analyses this work was supported by the state of Baden-Württemberg through bwHPC (MLS-WISO) and the German Research Foundation (DFG) through grant INST 35/1134-1 FUGG.

## Author Contributions

Designed the study: SLU, FG; supervised the project: SLU, FG; conducted experiments: KDC; analyzed data: KDC, SLU, PK, FG; developed the mathematical and computational models: PK, FG; performed the mathematical analyses: PK, TF, YS, FG; contributed important intellectual content: JH, HD; wrote the manuscript: PK, FG, SLU

## Supplemental Information

**Figure S1:**
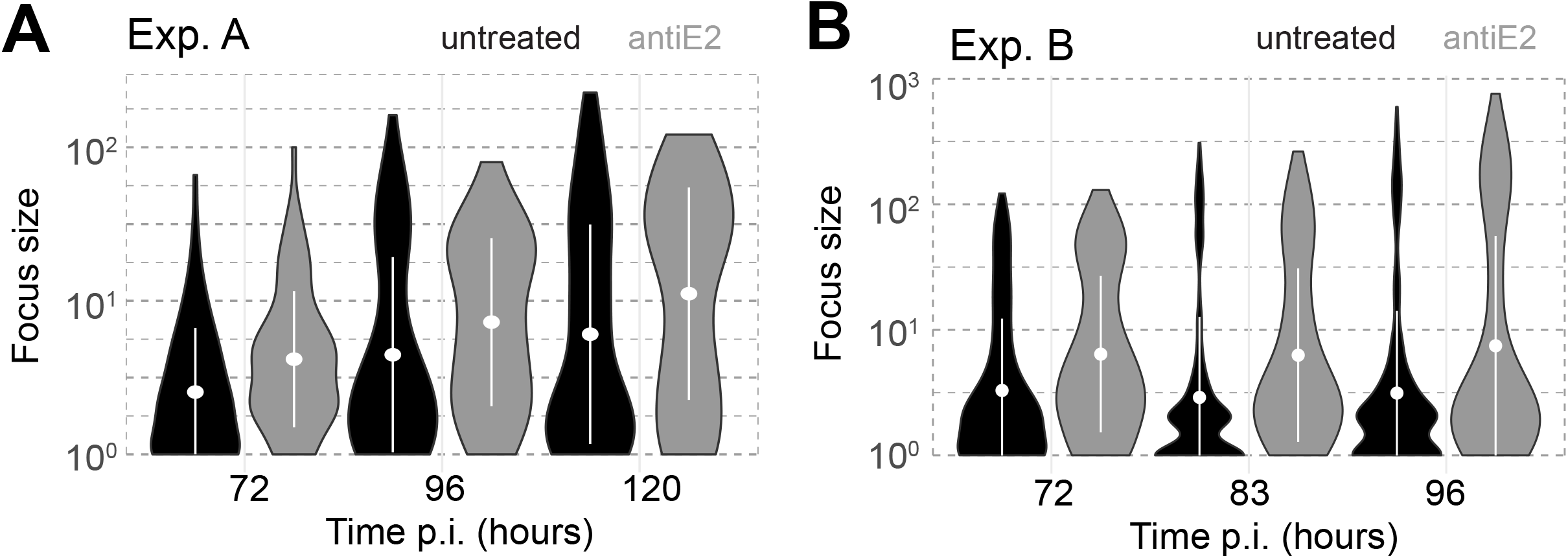
Bimodal HCV foci size distribution. Foci size distributions for Exp. A (**A**) and Exp. B (**B**) shown as violin plots corresponding to Figure 2B and C in the main text, respectively. Distributions are shown for untreated (black) and anti-E2 (grey) treated cultures with the mean (white dot) and standard deviation (white bar) indicated. The violin plots indicate the bi-modal distribution for anti-E2 treated cultures in Exp. B especially at later time points.

**Figure S2:**
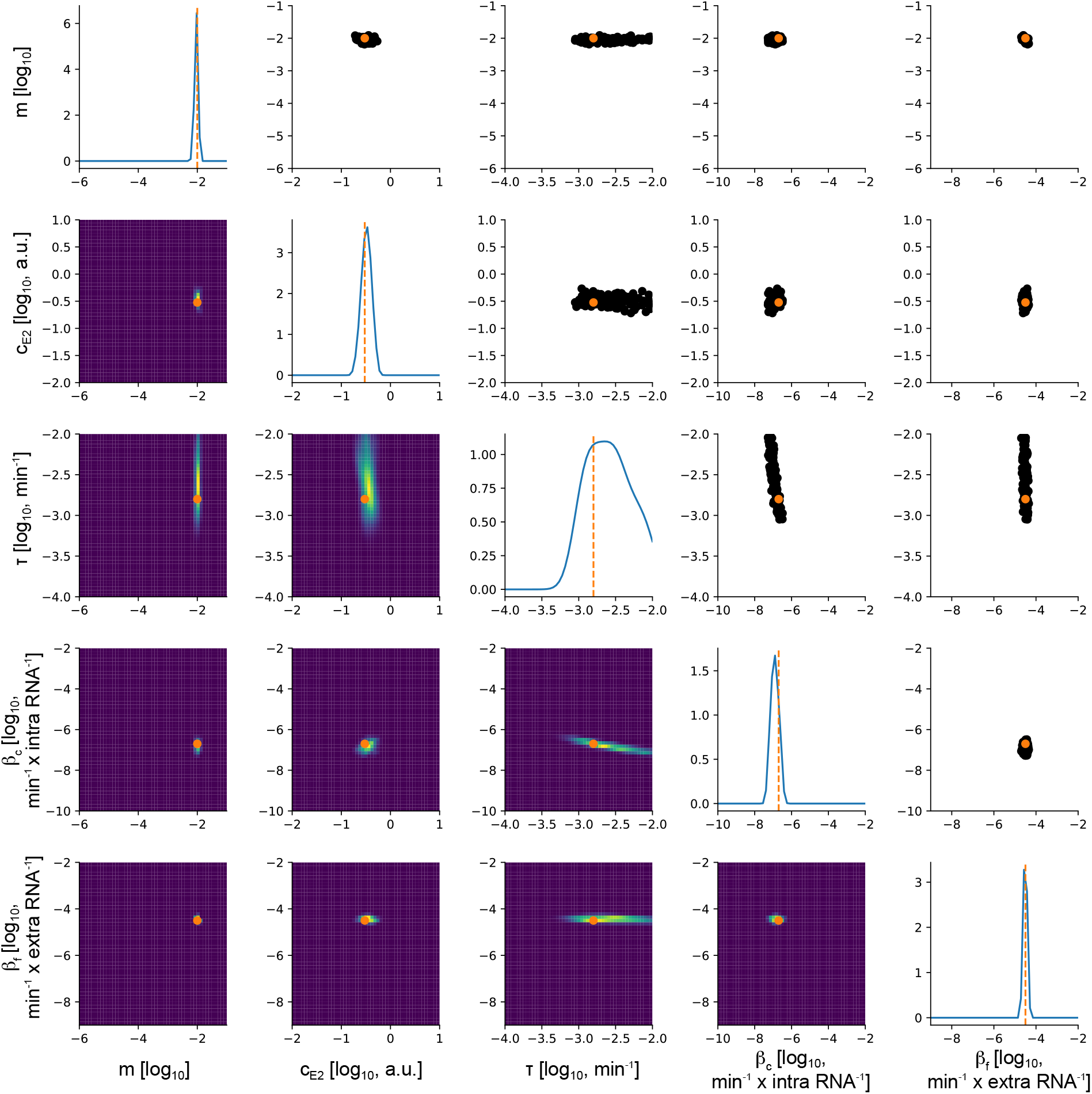
Estimated kernel densities of fitted parameters for *simulated* data. Each particle within pyABC used 5 individual ABM-simulations to calculate average infection dynamics according to the experimental conditions. Results shown were obtained after 17 pyABC generations. Individual rows and columns represent the effective diffusion rate of extracellular virus indicated by the coupling coefficient *m* (1^st^ row/column), the usage rate of anti-E2 *c_E2_* (2^nd^ row/column), the cell-to-cell infectivity parameter *τ* (3^rd^ row/column), and the cell-to-cell (4^th^ row/column) and cell-free (5^th^ row/column) scaling factor *β_c_* and *β_f_*, representing the corresponding transmission rates. Panels above the diagonal represent 100 individual parameter combinations (black), which had a distance smaller than 1.67 and below the diagonal the corresponding estimated two-dimensional kernel densities. The kernel density estimates of each parameter are shown on the diagonal. The parameters used for simulation are indicated in orange.

**Figure S3:**
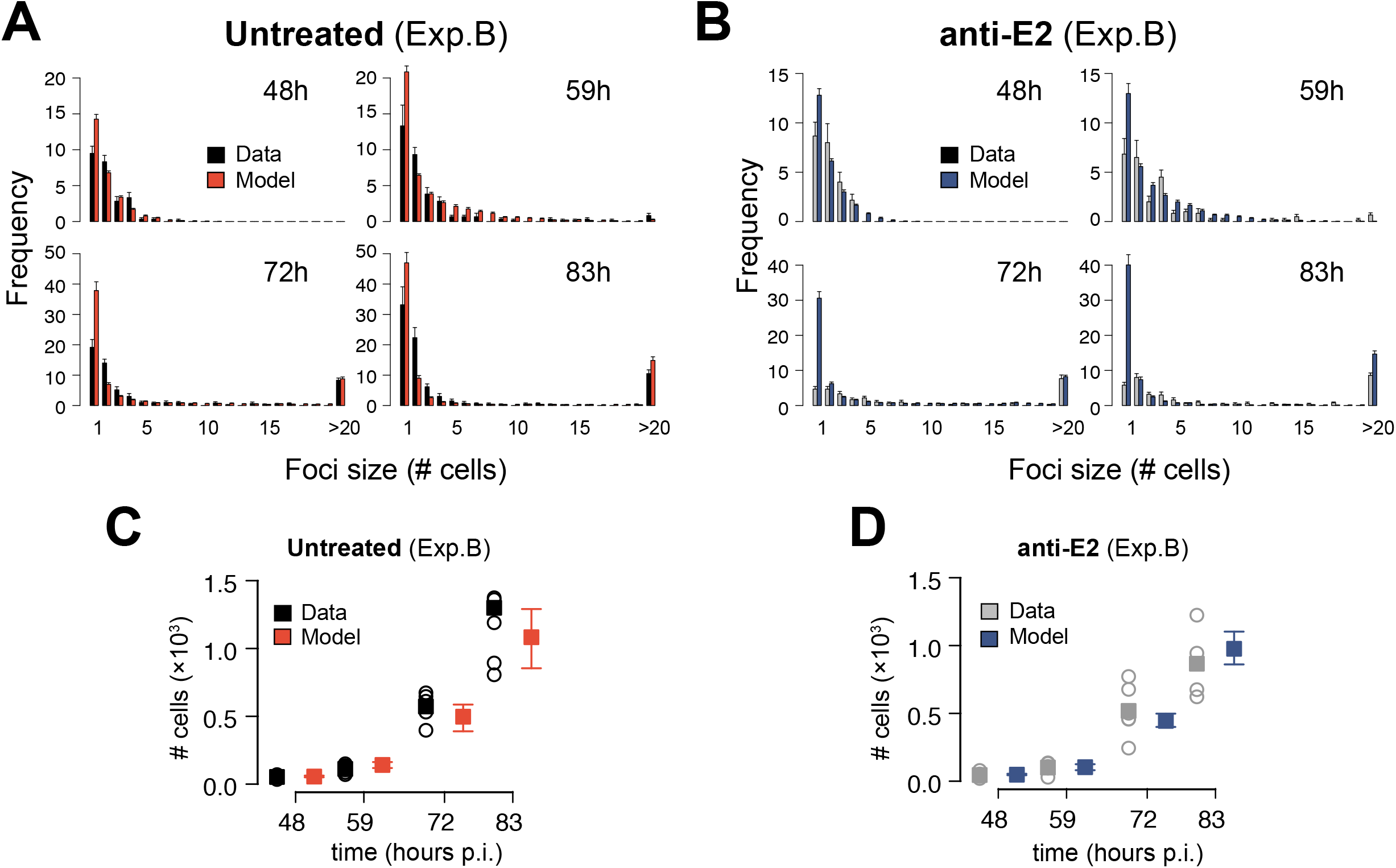
Model predictions for Exp. B: (**A,B**) Measured (black/grey) and predicted (red/blue) focus size distributions for Exp. B without (**A)** and with (**B**) administration of anti-E2 at 48, 59, 72 and 83 hours p.i. and calculated across 5 replicates according to the experimental conditions. Predictions are based on an exemplary parameterization of the model as obtained by the fitting procedure. **(C,D)** Predicted (red/blue) and measured (black/grey) average number of infected cells in untreated (**C**) and anti-E2 treated (**D**) wells calculated across 5-6 replicates.

**Figure S4:**
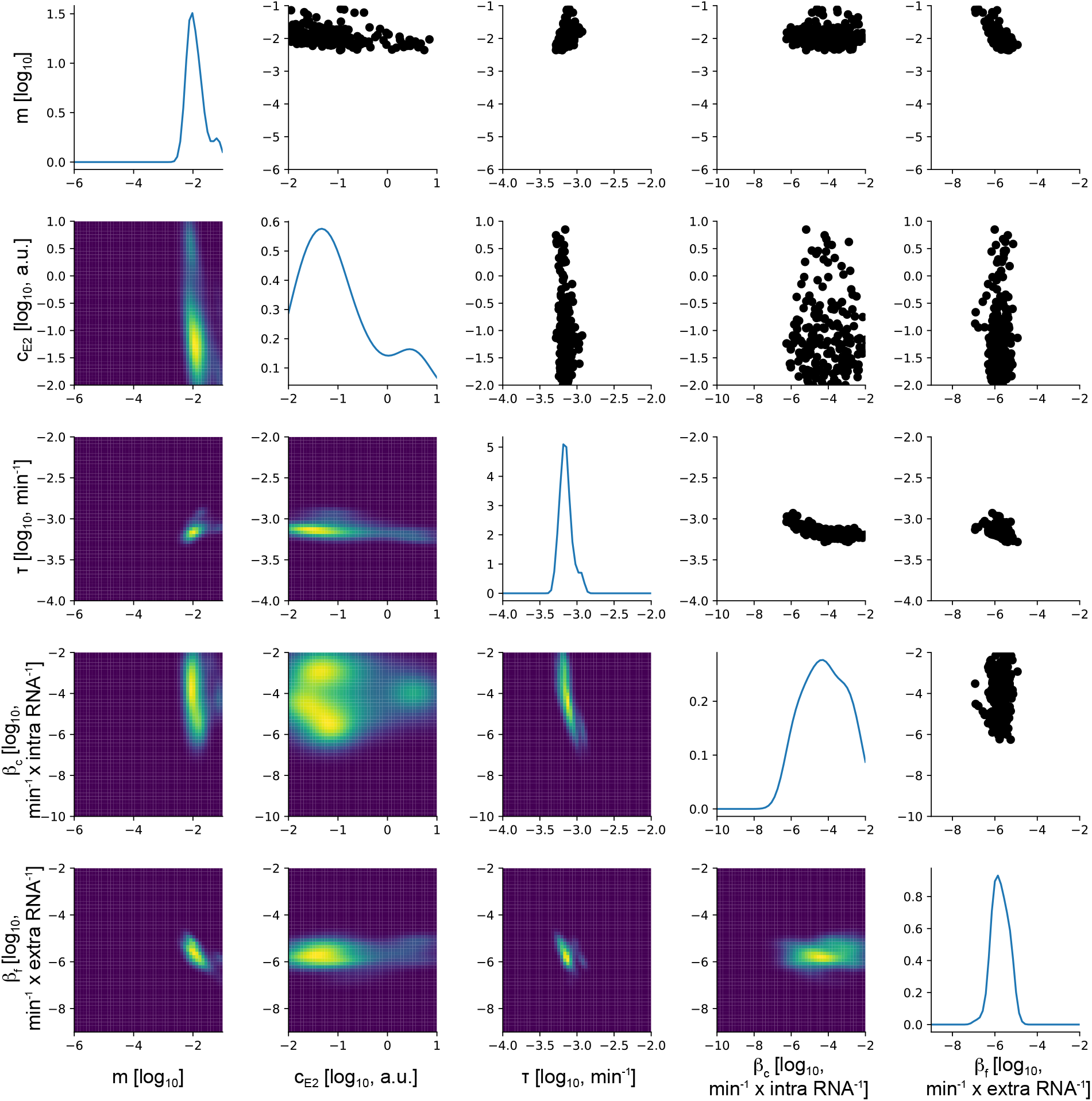
Estimated kernel densities of fitted parameters to *in vitro* data obtained from Exp. A. Each particle within pyABC used 2 individual ABM-simulations to calculate average infection dynamics according to the experimental conditions. Results shown were obtained after 15 pyABC generations. Individual rows and columns represent the effective diffusion rate of extracellular virus indicated by the coupling coefficient *m* (1^st^ row/column), the usage rate of anti-E2 *c_E2_* (2^nd^ row/column), the cell-to-cell infectivity parameter *τ* (3^rd^ row/column), and the cell-to-cell (4^th^ row/column) and cell-free (5^th^ row/column) scaling factor *β_c_* and *β_f_*, representing the corresponding transmission rates. Panels above the diagonal represent 200 individual parameter combinations (black), which had a distance smaller than 4.46 and below the diagonal the corresponding estimated two-dimensional kernel densities. The kernel density estimates of each parameter are shown on the diagonal.

**Figure S5:**
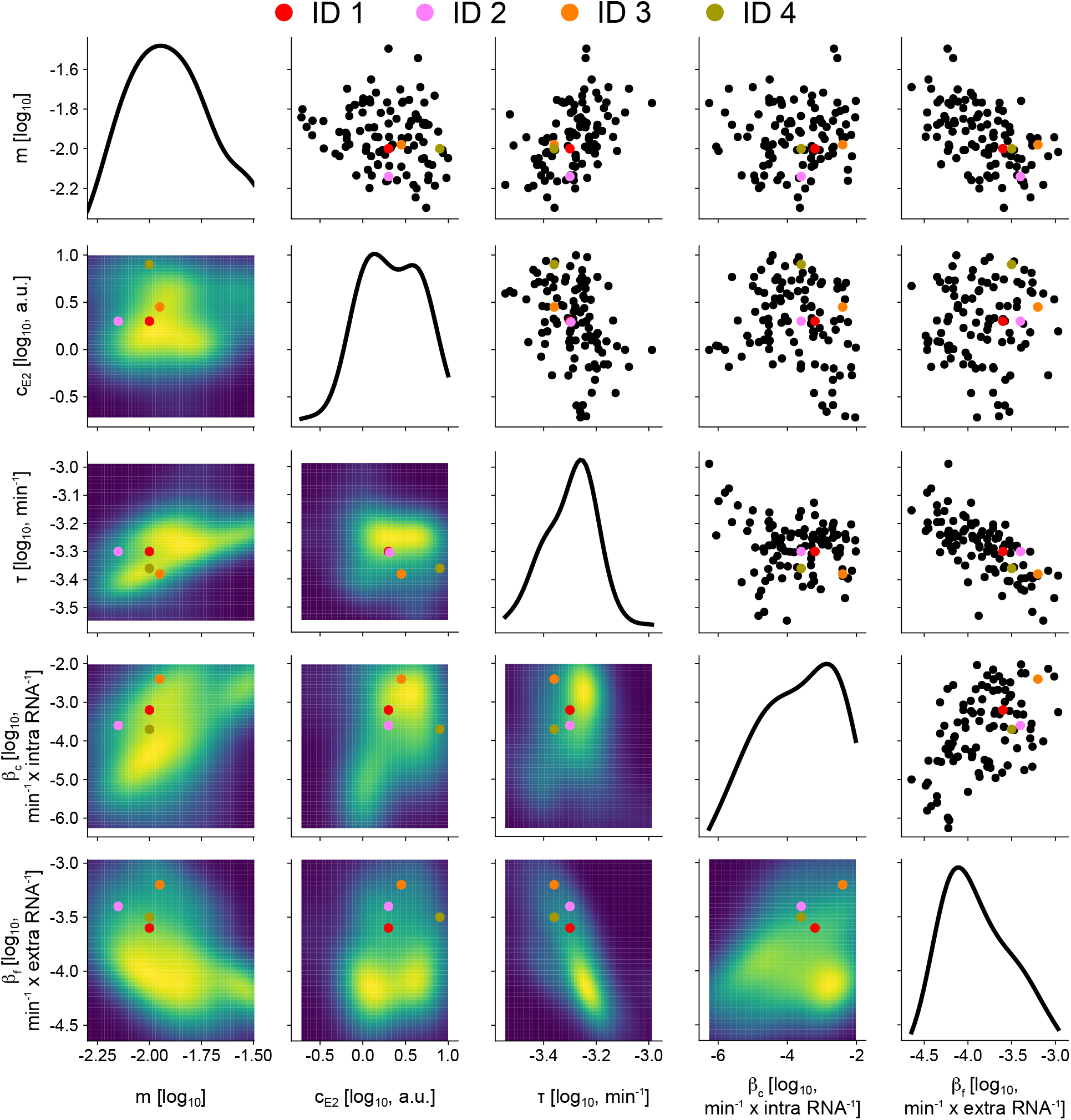
Estimated kernel densities of fitted parameters to *in vitro* data obtained from Exp. B. Each particle within pyABC used 5 individual ABM-simulations to calculate average infection dynamics according to the experimental conditions. Results shown were obtained after 13 pyABC generations. Individual rows and columns represent the effective diffusion rate of extracellular virus indicated by the coupling coefficient *m* (1^st^ row/column), the usage rate of anti-E2 *c_E2_* (2^nd^ row/column), the cell-to-cell infectivity parameter *τ* (3^rd^ row/column), and the cell-to-cell (4^th^ row/column) and cell-free (5^th^ row/column) scaling factor *β_c_* and *β_f_*, representing the corresponding transmission rates. Panels above the diagonal represent 100 individual parameter combinations (black), which had a distance smaller than 4.64 and below the diagonal the corresponding estimated two-dimensional kernel densities. The kernel density estimates of each parameter are shown on the diagonal. The parameters of the four best fitting particles after 13 generations of optimization with pyABC are indicated with red (ID 1), rose (ID 2), orange (ID 3) and olive green (ID 4).

**Figure S6:**
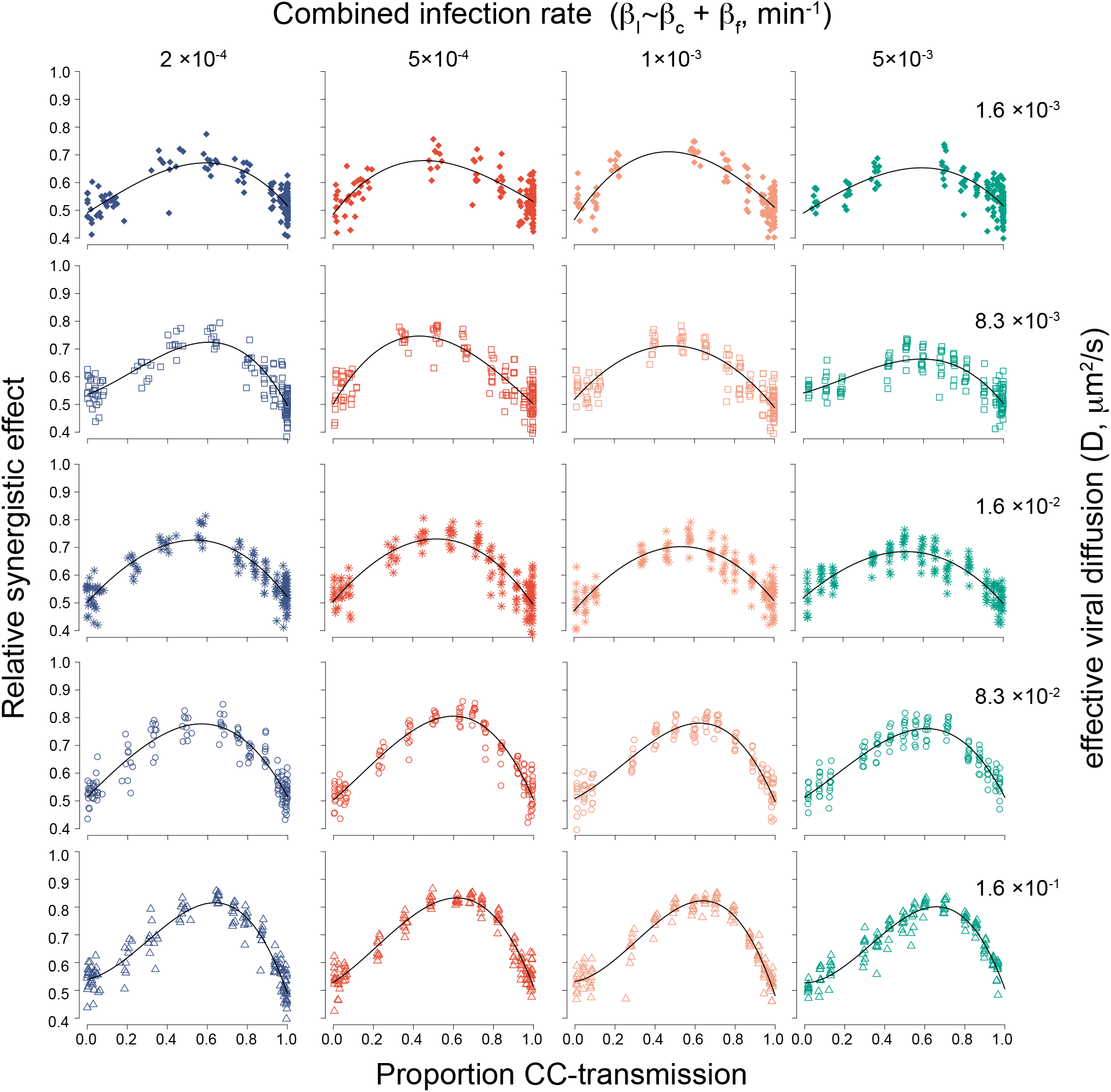
Advantages of combined modes of viral spread: Predicted advantage of combined modes of viral spread indicated by the Relative Synergistic eEfect (*RSE*) of combined viral spread (Expanded set of combinations compared to main manuscript **Fig. 5**). The *RSE* is shown dependent on the proportion of infections by cell-to-cell transmission for different assumptions of the combined occurrence of infection by both transmission modes and the viral diffusion coefficient. Points indicate the maximal *RSE* obtained for simulating viral spread in the ABM with different combinations of *β_c_* and *β_f_* making up the combined infection rate *β_I_*. Curves show the result of a polynomial of 3^rd^ degree fitted to the individual data.

**Figure S7:**
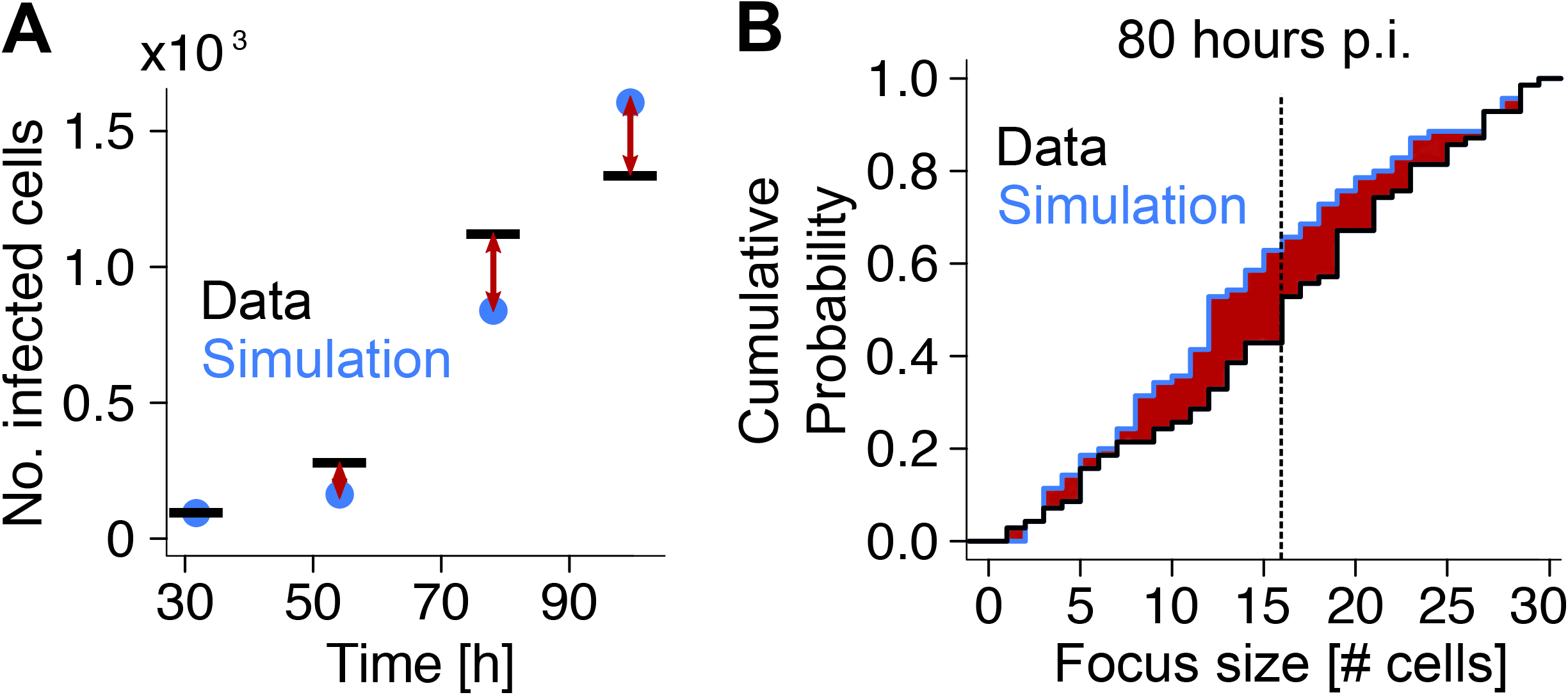
Graph of distance measure used for fitting ABM to *in silico* or *in vitro* data. **(A)** Shown is the absolute distance (red arrows) between hypothetical experiment (black lines) and prediction (blue dots). **(B)** The enclosed area of the hypothetical measured (black) and predicted (blue) cumulative density functions of the focus size distribution after 80 hours is indicated with the red shaded area. The dashed line indicates the measured average focus size.

---

**Table S1:** Parameter estimates for viral lifecycle kinetics using the mathematical model describing intracellular viral replication and viral export (see *Materials and Methods* for a detailed explanation of the individual parameters). Shown are the best fits with numbers in brackets indicating the 95%-confidence intervals obtained by profile likelihood analysis.

| Parameter | Description | Unit | Estimate [ 95%-CI] |
| --- | --- | --- | --- |
| $R_0$ | Initial intracellular HCV RNA | RNA cell<sup>-1</sup> | 80.1 [42.2, 192] |
| $V_0$ | Initial residual extracellular HCV RNA | $\times 10^3$ RNA cell<sup>-1</sup> | 5.91 [3.66, 10.5] |
| $\lambda$ | Intracellular RNA production rate | h<sup>-1</sup> | 0.24 [0.19, 0.28] |
| $\rho$ | Viral export rate | $\times 10^{-3}$ h<sup>-1</sup> | 2.10 [1.24, 3.47] |
| $R_c$ | Carrying capacity of intracellular RNA | $\times 10^4$ RNA cell<sup>-1</sup> | 7.73 [6.30, 9.76] |
| $c$ | Viral loss rate | $\times 10^{-2}$ h<sup>-1</sup> | 7.58 [4.88, 11.0] |

